# Quantified ensemble 3D surface features modeled as a window on centric diatom valve morphogenesis

**DOI:** 10.1101/468884

**Authors:** Janice L. Pappas

## Abstract

Morphological surface features are a record of genetic and developmental processes as well as environmental influences. The 3D geometric “terrain” of the surface consists of slopes via tangents, peaks and valleys via normals, smoothness of the transition between peaks and valleys, and point connections as flatness or curvature among all features. Such geometric quantities can be used to indicate morphological changes in valve formation over time. Quantified 3D surface features as geometric pattern ensembles may be representative of structural snapshots of the morphogenetic process.

For diatoms, valve formation and pattern morphogenesis has been modeled using Turing-like and other algorithmic techniques to mimic the way in which diatoms exhibit the highly diverse patterns on their valve surfaces. How the created surface features are related to one another is not necessarily determined via such methods. With the diatom valve face structure of layered areolae, cribra, and other morphological characters, valve formation exhibits different combined geometries unfolding as 3D structural ensembles in particular spatial arrangements. Quantifying ensemble 3D surface geometries is attainable via models devised using parametric 3D equations and extracting surface features via partial derivatives for slopes, peaks and valleys, smoothness, and flatness as feature connectedness. Differences in ensemble 3D surface features may be used to assess structural differences among selected diatom genera as indicators of different valve formation sequences in surface generation and morphogenesis.

## Introduction

As a guiding force historically in biology, the interweaving of evolution and genetics comprise the modern synthesis (e.g., Huxley 1942; Simpson 1946) and the advent of mathematical population genetics showed that the application of quantitative methods was efficacious in understanding the role of genetics in evolution (e.g., Fisher 1918). To account for advances in genomics and molecular biology as well as developmental theory and morphology, and including abrupt timing events such as endosymbiosis (e.g., Kutschera and Niklas 2004), the extended evolutionary synthesis (e.g., Pigliucci et al. 2006) and post-modern evolutionary synthesis (e.g., Koonin 2009) propose to remedy the shortcomings of the modern synthesis. Macroevolution across taxonomic levels going beyond species-specific biology is also a contributing factor in the impetus for a more accurate synthesis in understanding the biological world. The phenotype and its role in morphological evolution and evolutionary developmental biology (e.g., Gilbert 2003; Organ et al. 2015) are of particular interest in studying morphogenesis from a macroevolutionary viewpoint.

Developmental processes encompass information transfer over time. Some of the information transfer involves the phenotype of the organisms as changes in morphology and evidenced in morphological variation. The dynamics of form and structure comprising morphogenesis as an evolutionary process occurs via various mechanisms and at various spatial and temporal scales. Constraints on the kinds of phenotypes that evolve may be a result of the morphogenetic process, producing specific outcomes evidenced more broadly as morphological evolution. Morphogenesis is expressed externally in the morphological patterns of organisms as a result of internally occurring genetic, cytological and developmental processes. Through developmental processes, morphogenetic outcomes may be the result of further refined evolutionary avenues so that such developmental constraints may produce morphological variation more specifically among taxa as phenotypic expression via epistasis and canalization (e.g., Waddington 1942).

Phenotype during development may be studied in unicellular organisms with regard to epistasis (e.g., Sanjuán and Elena 2006) and canalization (e.g., Yadav et al. 2016). The processes of epistasis and canalization concerning the genotype may be extended to understand interactions among quantified characters of the phenotype in models of development (Rice 1998). Phenotypic characters are representations of morphological features, and their quantification enables a way to model inferences of their relation to information transfer during the life cycle and development of a unicellular organism.

As changes in phenotype occur, morphogenesis during unicell development may be quantifiable as 3D surface features. Ensembles of these surface features may be used to track changes in phenotype over time using morphological 3D models. Documenting these phenotypic changes via 3D surface models enables studying the development of a whole unicellular organism more comprehensively in 3D rather than just looking at morphology as isolated or individual parts with no reasonable way to provide a cohesive view of the entire unicellular organism.

### From 3D surface morphology to morphogenesis

Morphology of an organism is what we perceive. We look at an organism to register its external features in our mind—its size, its color, its shape. When describing these features, we often characterize them as measurable and countable attributes with boundaries. That is, we assign numerical descriptors about how much we see. For example, the length of a given feature, the number of repetitions of a given feature, and the amount of color and geometry of shape of a given feature are all observations that tell us something measurable about morphological attributes. All of these morphological measures are interpreted to have biological meaning.

Other aspects of morphology are evident, but not necessarily so easily summarized as a countable feature. How much does the surface slope? How do peaks and valleys vary on the surface? How smooth is the surface? Are some places on the surface much more pronounced than others? How flat or curved is the surface? How are surface attributes connected to one another? All of these morphological attributes may have biologically significant meaning that is just as important as countable features. Such surface features may be important in the dynamic growth of an organism and indicative of structural patterning. Morphological changes during growth potentially signifying phenotypic plasticity in which repeating patterned structures and their variation may be recorded. Such variation may be seen at the species or higher taxonomic levels via non-countable morphological surface features.

Morphological surface features are a record of the processes necessary to produce the way an organism looks and are identifiable from surface points to combinations of geometric arrangements of surface points and contours. The geometric “terrain” of the surface consists of slopes via tangents, the peaks and valleys via normals, smoothness at transitional zones between peaks and valleys, and point connections as curvature or flatness between tangents and normals spatially tying multiple features together. For diatoms, surface features may be functionally important in predation resistance or defense (peakedness), glide ability in water (smoothness), ability to be cloaked or disappear into the environment (flatness), or countershading and reflectability in the environment (slopeness). Quantified surface features may be used to represent structural features and changes during morphogenesis as ensembles. That is, the geometry of morphology is quantifiable as ensemble surface features in organisms. Because of their elaborate geometric morphology, diatoms would qualify as model systems (De Tommasi et al. 2017) for study using ensemble surface features (Fig. 1).

**Fig. 1.**
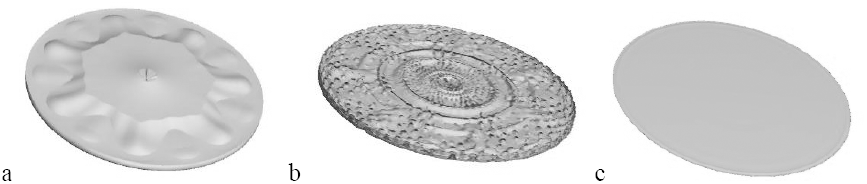
Ensemble surface features--examples: a, smooth, but not flat; b, flat, but not smooth; c, smooth and flat. Smoothness is determined from the Laplacian, and flatness is determined from Christoffel symbols.

For diatoms, silica valve formation and pattern morphogenesis has been of interest and modeled using Turing (1952)-like and similar algorithmic techniques (e.g., Gordon and Drum 1964; Parkinson et al. 1999; Gordon et al. 2009). Models have been created to mimic the way in which diatoms exhibit the highly diverse patterns evident on their valve surfaces. Techniques including artificial neural networks via multi-agent algorithms or cellular automata systems have been used in various attempts to represent “mechanistically” the way the biological world works in all its complexity. Results from these techniques and algorithms have an intuitive appeal because of their resemblance to the consequents of biological processes. It is tempting to think of such algorithms as “biological mechanisms” when, in fact, they are not. They are only pattern generators. To date, such techniques have been useful in creating patterns but have not been integrated and matched at multiple scales with actual silica deposition events, developmental or life cycle changing events, or any link to biologically-evidenced morphogenetic steps in diatoms. Initiation and termination events of techniques such as those generating reaction-diffusion patterns, if they do occur in diatom valve formation, has yet to be addressed in more than an ad hoc way. Different applications of reactive-diffusion as different models may have multiple mechanistic explanations or make incongruent predictions (e.g., Cotterell et al. 2015). Despite the recent renewed interest in reaction-diffusion systems at the molecular and nanoscale levels (e.g., Green and Sharpe 2015), diatom valve patterning is a 3D process occurring during morphogenesis. Caution must be exercised concerning pattern-generating models as being equivalent to actual biological processes (Wimsatt 1987).

Having said this, the utility of modeling has been demonstrated time and time again as an aid to gain an understanding of the potential explanations for biological processes (e.g., Laubichler and Müller 2007). Vegetative diatom cell morphology as a geometric construct can be used in a phenomenological approach and basis for modeling. From the whole organism cell pattern, the mature diatom vegetative cell is represented by valve face geometry, and changes in morphology during valve formation may be achieved via back-tracking from this cell to the start of silica deposition. Organism surface represents the boundaries of its three-dimensional (3D) morphology and as such, this surface geometry may be used as a proxy for the phenotype (e.g., Pappas and Miller 2013). Considering surfaces may enable a better understanding of potential function and evolution of biological structures and requires looking at 3D surface geometry and ensembles of those geometries.

For diatom valve formation, 3D surface geometry is necessary to consider the whole cell. Morphological changes over time occur in 3D, and all the information available on the surface is of importance concerning morphogenesis. Although quantitative techniques via a closed curve orthogonal polynomial or matching arbitrary boundary points among species-specific taxa via deformation techniques have been used to characterize valve shape in diatom identification and classification studies in taxonomic analysis (e.g., Pappas et al. 2014), these techniques are inadequate in the assessment of surface morphology. Circular and oval centric valve outlines provide no differentiating information because of their shape. Even if differentiation of outline shapes is possible, this says nothing about the information contained in and on the valve surface. For diatom valve morphogenesis, 3D quantitative methods enable the inclusion of all boundary points on the 3D surface.

Diatom valve formation during morphogenesis may be “cataloged” as geometric surface changes. Absent direct matching with biological processes, a geometric interpretation can be qualitatively relational in terms of ensembles of morphological changes over time. With the striking geometry of the scaffolded structure of the diatom valve face of layered areolae, cribra, and other morphological characters, valve formation has different combined geometries from layer to layer, combining such structural elements in a spatial arrangement. In actual biological cells, the layering effect has been used as a means to examine structural morphology with imaging and microscopy techniques via *z*-stacking (e.g., Shihavuddin et al. 2017). Geometrically, a cell can be crudely thought of as a series of circles stacked one upon the other to form the 3D cylindrical shape. Its 3D reconstructed image is an analytic approximation of a real entity. Removal of each circular layer from a 3D cylinder would expose the relief of successive surfaces of a cell such as a centric diatom valve face. Back-tracking to the start of silica deposition, the whole diatom 3D model and its forming valves during morphogenesis may be devised as the reverse stepwise layering of successive surfaces of *z*-stacking. Layered additions as a sequence of steps may be modeled with *z*-layers to produce the whole cell. A 3D surface model is an implicit approximation of a real entity.

### Geometric basis of 3D surface models and analysis

Initially, a 3D surface model is created using parametric 3D equations. Geometric analysis of a 3D surface model necessarily means defining that surface as a tangent space. This space contains points in which first partial derivatives from the system of parametric 3D equations form the Jacobian matrix (i.e., the Jacobian) and represents the sloping of the surface. Connectivity of points on the surface necessitates the creation of ensembles of surface features because partial differentiation is insufficient in constituting connection of neighboring tangent spaces. To achieve connection of tangent spaces, Christoffel symbols are calculated as connectivity representing flatness or global curvature among points on the surface. In a tangent space, change in basis vectors relative to a basis forms the way in which to measure a differential change in coordinates in each coordinate direction (Koop 1993) on the surface, and change in coordinates occurs via the Jacobian (Kaplan 2003). Connection coefficients are Christoffel symbols in which the basis vectors change linearly. As a result, Christoffel symbols are a linear connection among ensemble surface features as they give structure to a tangent space.

The second partial derivatives are the Hessian matrix (i.e., Hessian) and Laplacian (i.e., Laplace operator expressed via Laplace’s equation) that are used to quantify the degree of local curvature of critical points and mapping of a one ensemble of surface features to another one on the surface, respectively. That is, the derivative of the slope of the tangent as peaks, valleys and saddle points (i.e., saddles) express maxima, minima and points of inflection on the surface, and the sum of the change among ensembles of these surface features express the degree of smoothness on the surface.

The Jacobian, Christoffel symbols, the Hessian, and the Laplacian express ensemble surface features as morphogenetic descriptors of each step of a diatom valve formation sequence as slopeness, connectivity via curvature or flatness, peak-to-valley changes, and smoothness, respectively, of the 3D surface. These quantities will characterize the valve surface as pointwise connectivity via degree of curvature or flatness and smoothness between elevations and depressions and sloping on a diatom valve surface.

### 3D surface analysis

#### Differential geometry of 3D surfaces

Each point on the surface is composed of a trihedron of vectors defined as a moving reference frame. For a curve, a Serret-Frenet frame consists of tangent (**t**), unit normal (**n**), and binormal (**b**) vectors, while for a surface, a Darboux frame consists of tangent (**t**), unit normal (principal normal) (**u**), and tangent normal vectors (**v**). At the tangent and tangent normal vectors of a point, normal planes at the unit normal can be used to cut the surface and define principal directions. Curvature, *κ,* at a point in relation to **n** and **u** is *κ***n** · **u** (Kaplan 1999).

Curvature is the rate of change of a tangent line to a curve (do Carmo 1976). The maximum and minimum curvatures at a point are the principal curvatures, *κ*_1_ and *κ*_2_. The product of the maximum and minimum curvatures is Gaussian curvature. Mean curvature is half the sum of the maximum and minimum curvatures. If both *κ*_1_ and *κ*_2_ are positive or negative, then the surface is locally convex. If *κ*_1_ is positive and *κ*_2_ is negative or vice versa, then the surface is locally a saddle. If *κ*_1_ or *κ*_2_ is equal to 0, then the surface is in between a convex and saddle. If *κ*_1_ and *κ*_2_ are both equal to 0, then the surface is a plane or a monkey saddle at a flat umbilic (Kaplan 1999).

Rate of change of a tangent is a first partial derivative with respect to a given curve on a surface with arc length, *s*. All of the tangent lines at every point on the surface are summarized in a Jacobian. Numerical solution to the Jacobian characterizes all the tangent lines and planes on the whole organism surface. Tangent lines define an intrinsic metric via the arc length and are coefficients of the first fundamental form, which is an inner product on the tangent space of a surface. The first fundamental form enables the measurement of length, angle, curvature, surface area and any other metric on a surface in 3D space. The second fundamental form defines tangent planes in terms of change in surface shape of the normals with respect to the tangents (Koenderink 1990) and is a quadratic form on the tangent space of a surface in 3D space. That is, the second fundamental form is a change of local shape over the surface.

For a two-dimensional (2D) patch on a 3D surface, tangent vectors span the tangent plane and serve as a basis in a matrix of coefficients for the differential of the tangent vectors. The matrix expressed in terms of the first and second fundamental forms represents Gaussian and mean curvature, respectively (do Carmo 1976). Using Gaussian curvature, shape can be determined as a general characterization of the surface (Koenderink 1990).

There are normals to tangent vectors spanning a tangent plane. The differentiable field of the unit normal vectors on the surface is a Gauss map of the tangent space of a point. The differential of a Gauss map is expressed as the shape operator or a Weingarten map. The shape operator determines all the tangent planes in the neighborhood of a point and is the change in surface normals with tangent vectors (Koenderink 1990).

Rate of change of a tangent line to a curve may be characterized by Gaussian curvature, *K* = *κ*_1_*κ*_2_, and mean curvature, *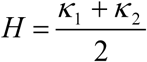*. Gaussian curvature is an intrinsic property of a 3D surface object without reference to the geometric space in which the object resides, and mean curvature refers to the embedding of a 3D surface object in a 3D space and is its extrinsic property (Koenderink 1990).

The first and second fundamental forms as well as Gaussian and mean curvature, respectively provides a link from Jacobian to Gauss and Weingarten maps (do Carmo 1976). The Jacobian determinant of a Gauss map is Gaussian curvature. Principal curvatures of a Weingarten map are eigenvalues and are an indicator of how much the surface bends (Koenderink 1990). Principal directions are eigenvectors of a Weingarten map, and from the second fundamental form, rotation of the *x, y* tangent plane produces eigenvectors of the Hessian (do Carmo 1976). By projecting the covariant derivative of the Weingarten map onto the second fundamental form, the connection gradient, ∇, is obtained, which is also the derivative of the trihedron of vectors and represented by Christoffel symbols (do Carmo 1976). The connection matrix of Christoffel symbols represents the connection of points and the movement of a Darboux frame as it relates to a Serret-Frenet frame on a curve (i.e., trihedron of vectors) on the surface (Koenderink 1990). The Laplacian depends on connectivity via the divergence of the gradient on the surface and is the trace of the Hessian (do Carmo 1976). The Jacobian is related to the Christoffel symbols because each row of the Jacobian is a gradient (do Carmo 1976; Kaplan 2003), and the Jacobian determinant relates surface connections via coordinate transformation (Sochi 2016). Gauss and Weingarten maps implicitly describe the functions that are used in characterizing 3D surface morphology.

#### 3D surface feature geometry and morphological attributes

Changes in the contours of the 3D surface or steepness or shallowness of slopes at a given *u, v* coordinate can be characterized in the *x*-, *y*-, and *z*-directions resulting in a Jacobian. The steepest and shallowest points as well as those points that indicate a transition from shallow to steep slopes (or vice versa) are calculated as second partial derivatives. These slopes are quantifications of local maxima, minima and saddle points (i.e., saddles) with the highest maximum and lowest minimum as global values. The Hessian consists of second partial and mixed partial derivatives so that the Jacobian is transformed from a six element to a 12-element matrix.

When considering a patch as a tangent plane, the Hessian is much easier to calculate. Because *x* and *y* define a plane, only the *z*-direction needs to be considered, and in this case, the patch is a Monge patch (Koenderink 1990). The *z*-direction is the height at each point on a patch, and the partial derivatives of *z* with respect to *u* and *v* from the Jacobian are the maximum and minimum slopes that represent a morphology gradient of the direction of maximum height of the surface as normals to the tangent plane of the surface. The magnitude of maximum height is the square root of the sum of the second partial derivatives of *z* with respect to *u* and *v*. These second partial derivatives along with the mixed second partial derivatives comprise the Hessian for *z*. Mixed second partial derivatives of *z* represent a combination of slopes of the morphology at that point on the surface. Movement of the change in height along a curve on the surface determines the effect of a combination of slopes on the characterization of morphology.

From the Hessian, the sum of the elements of the diagonal is the Laplacian, ∇^2^, and is the divergence of the gradient on the surface. Taylor expansion of the Laplacian around zero gives the state of the surface under conditions of no change in equilibrium and as a measure of stability that is solvable by finite differences (Weinberger 1965). Stability at equilibrium is also equivalent to being a measure of smoothness. Each of the terms from the finite differences approach with respect to *u* and *v* are eigenvectors, and the first term may be used to indicate approximate smoothness. Because the diagonal elements of the Hessian are used to calculate the Laplacian, it is these elements that are the eigenvalues of the eigenvectors. The variation in smoothness is evaluated by the sign of the sum of the eigenvalues or trace that is the Laplacian. That is, the Laplacian measures the degree to which perturbations on the surface reflect the smoothness of surface morphology.

Surface feature connections via Christoffel symbols are a measure of the degree of flatness or curvature of the surface (Koop 1993). For Christoffel symbols that are vanishing, the surface is flat. That is, for zero Gaussian curvature and Christoffel symbols at zero, the surface is flat (Misner 1973; Sochi 2016). The connection between highly textured and smooth surfaces are summarized in the Christoffel symbols as an indicator of changing curvature and flatness both vertically and horizontally across a contiguous surface morphology.

### Centric diatoms and valve formation in morphogenesis

#### Taxa used as exemplars in 3D surface models in diatom morphogenesis

Centric diatom taxa chosen for study as exemplars were *Actinoptychus senarius, Arachnoidiscus ehrenbergii* and *Cyclotella meneghiniana*. These taxa were readily modeled using parametric 3D equations and represent three widely different valve surfaces exhibiting distinctly different features associated with silica deposition during valve formation. Ensemble valve surface features are readily measured to illustrate quantitatively the distinctions among the taxa during valve formation as a part of the morphogenetic process.

Of the three exemplar centric diatom taxa, *Actinoptychus senarius* exhibits a circular buckling pattern as alternating undulations of sectors emanating from the center (Gordon and Tiffany 2011). Such a pattern may be recoverable as a wave front (Nechaev 2017). Rimoportulae are curled, and the valve is divided into sectors by interior folds or walls (Hasle and Sims 1986; Lee and Chang 1996). For *Actinoptychus senarius*, the areolae are a coarse or reticulate pattern on the valve surface, and there is a marginal ridge from which external projections of the rimoportulae protrude above the valve face ((Hasle and Sims 1986). *Actinoptychus senarius* has a narrow hyaline zone that extends from the central area to around halfway to the valve margin. Heterovalvy is not present in *Actinoptychus senarius* (Gordon and Tiffany 2011).

*Arachnoidiscus ehrenbergii* may exhibit heterovalvy during auxospore attachment to the epivalve of the parent cell (Kobayashi et al. 2000; Sato et al. 2004). Different valve patterns may form as a result of differing attachment sites. Re-creation of a diatom valve was accomplished using digital holography, resulting in a model of *Arachnoidiscus* that was obtained via image reconstruction of an optical wave front (Ferrara et al. 2014; Ferrara et al. 2016). The model showed elongated central slits in a ring surrounding a small planar central area on the valve and pores diminishing in view of a planar field (Ferrara et al. 2014). The ribs emanating from the central area are regularly spaced and may be bisected to look like windows on the pore structured pattern below (Brown 1933).

*Cyclotella meneghiniana* has distinctive striations as ribs regularly placed at the valve margin covering about half of the valve face (Schmid and Volcani 1983; Håkansson and Chepurnov 1999). These striations form a regular undulating pattern around the valve periphery giving *Cyclotella meneghiniana* its distinct appearance. Marginal rimoportulae are present as well (Håkansson and Chepurnov 1999), and the central area has fultoportulae. The clear central area exhibits a rather uniform surface (Schmid and Volcani 1983; Håkansson and Chepurnov 1999). *Cyclotella meneghiniana* may exhibit heterovalvy (Round et al. 1990).

#### Morphogenetic descriptors of centric diatoms in valve formation as sequential change in 3D surface morphology

Diatom growth patterns follow two general tracks—one for centric diatoms, and the other for pennate diatoms (Round et al. 1990). Centric diatom valve formation may be characterized simply as growth from an annulus to the valve margin, while pennate diatoms exhibit growth from a sternum to the valve margin (e.g., Schmid and Volcani 1983). Growth is horizontal across the valve face in 2D and horizontal and vertical in 3D. In the vertical aspect of the diatom growth system, the 3D surface may be characterized as layers of morphology. Quantitatively, as each layer is accounted for, vertical growth of the organism can be modeled using a series of systems of parametric 3D equations per taxon. The relation between developmental growth and 3D morphology is measurable as the change in whole surface morphology by additional surface morphologies, i.e., ensemble 3D surface morphologies. This change is more precisely a starting morphology plus an additional morphology. The addition is integrated into the original whole so that a new system of parametric 3D equations emerges from the original system. As subsequent additions are made, the cumulative result is modeled as the final valve produced after valve formation is completed.

Schmid and Volcani (1983) determined that diatom valve formation roughly occurs as a three-stage process. For centric diatoms, the starting point is the annulus from which radial rows of silica are deposited in a horizontal fashion toward the valve margin in a crosswise pattern with connections so that a branching pattern materializes. This is the basal layer in which silica is deposited to initiate the rudimentary formation of internal rimoportulae as well, from which the next stage of valve formation occurs. This next stage involves the initiation of vertical silica deposition where the walls of the areolae increasingly emerge, and the round pores that may be formed in the first stage become hexagonal or more geometrically defined in shape as well as the formation of external rimoportulae occurring. Cribra and cribella mark their formation during the third stage in which completion of horizontal silica deposition occurs. The size and spacing of these structures are controlled by areolae (Schmid and Volcani 1983) as the successive layers of silica are deposited to complete the valve structure to the margin (Rogerson et al. 1986).

Within the silicalemma and silica deposition vesicle (SDV), new wall formation occurs as silica is aggregated (Schmid and Volcani 1983; Rogerson et al. 1986; Round et al. 1990), and the plasmalemma molds cell wall shape prior to valve formation (Schmid 1987). From the center to the valve margin, structural changes occur in a sequence over time. Common valve structures are created for a given species within a genus, and differences in those structures occur via silica deposition for multiple species within that genus in terms of the location and timing of structural elements emerging during valve formation.

### Purposes of this study

In this study, Schmid and Volcani’s (1983) schema is used to characterize ensemble 3D surface changes in the modeling of centric diatom valve formation stages during diatom morphogenesis. Models will be created using systems of parametric 3D equations for each sequence of valve formation steps. The ensemble 3D geometric surface properties are quantified as the Jacobian, Hessian, Laplacian, and Christoffel symbols from slopes, peaks-valleys-saddles, smoothness, and connectedness via flatness or curvature, respectively. Ensemble surface geometric features and the degree to which they change during valve formation is used to characterize a constructed diatom morphogenetic system with regard to silica deposition implicitly over time. Implications of ensemble surface features used in characterizing a process such as morphogenesis are explored for macroevolutionary processes such as epistasis and canalization.

## Methods

Diatom valve formation is depicted using 3D surface models representing vegetative cells in 3D (*x, y, z*) space. Parametric 3D equations are interpretable in terms of their utility in relating 3D surface geometry to 3D surface morphology. Variables *x, y, z* parameterized by *u, v* are used in the characterization of a point anywhere on the surface as well as movement from point to point on the surface along a given curve on a given diatom model. Systems of parametric 3D equations of the general form *x* = *f*(*u, v*), *y* = *g*(*u, v*), *z* = *h*(*u, v*) with Cartesian coordinates *x, y, z* in parameters *u, v* and evaluated on the interval [0, 2*π*] are used to create the 3D surface models (Pappas 2005a, b; Pappas 2008; Pappas 2011; Pappas and Miller 2013; Pappas 2016). Parametric 3D equations enable the quantification of points on curves and/or surfaces so that each point in *x, y, z* Euclidean 3D space can be mapped to a new 2D *u, v* Euclidean space. Because surface morphology resides within a curved boundary (e.g., circle, oval or other polygonal 2D form), the new 2D space contains morphological information in *u, v* surface points. Each 2D surface point is a base for a vector in which the arrowhead of the vector points away from the surface. Using vectors that point perpendicularly from the surface at each point, maximum changes from point to point are recorded changes as one travels along the surface in any direction. The *u, v* parameters are 2D points extracted from a 3D *x, y, z* diatom valve surface model that contain information on changes in the topography of the surface. These changes are measurable as the solution of parametric 3D equations that map the *x, y, z* variables to the *u, v* parameters. In this way, 3D surface geometry is used in quantifying 3D surface morphology.

### Measurement of ensemble surface features and 3D surface morphology: derivation and solution of the Jacobian, Hessian, Laplacian, and Christoffel symbols

#### The Jacobian of 3D surface morphology

From parametric 3D equations, first derivatives are expressed in terms of first partial derivatives with respect to arc length, *s*. The differential of *s* is *ds*^2^ = *dx*^2^ + *dy*^2^ + *dz*^2^ on the surface, and 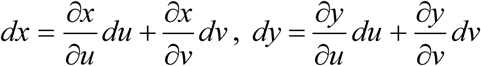, and 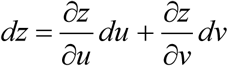. The Jacobian is 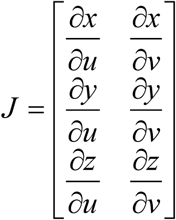 as first partial derivative of *x, y, z* with respect to *u, v* (Kaplan 2003). Numerical solution to the Jacobian characterizes all the tangent lines and planes on the whole organism surface. Eigenvalues, *Ψ*, of the Jacobian may be negative, meaning a stable surface at extrema (i.e., maxima or minima), or positive, meaning an unstable surface at extrema (i.e., saddles or other extrema that are neither maxima nor minima).

#### Monge patch

For a whole organism 3D surface expressed parametrically as *x* = *f* (*u, v*), *y* = *g*(*u, v*),*z* = *h*(*u, v*), *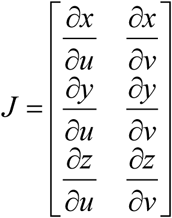*. For a patch on or as that surface, *x* = *g*(*u, v*), *y* = *h*(*u, v*), *z* = *f* (*x, y*) = *f* [*g*(*u, v*), *h*(*u, v*)], and 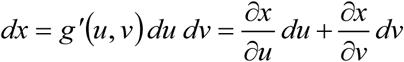, 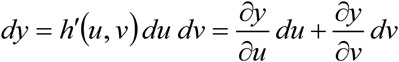,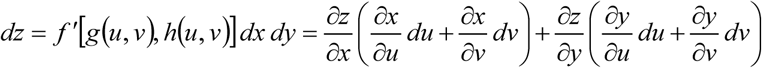. For *J*, define each row as a linear combination of the other. At (*u*_0_, *v*_0_), the submatrices are 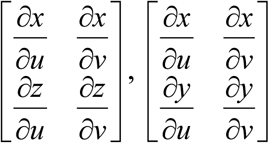 and 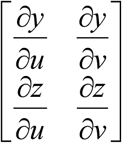 for (*x, z*) = (*x*(*u, v*), *z*(*u, v*)), (*x, y*) = (*x*(*u, v*), *y*(*u, v*)), and (*y, z*) = (*y*(*u, v*), *z*(*u, v*)), respectively. By the inverse function theorem, the reparameterizations to Monge patches are 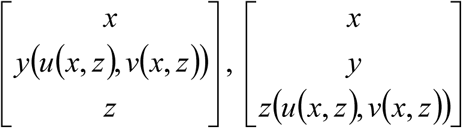, and 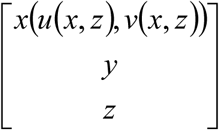, respectively.

#### First and second fundamental forms and surface characterization of the Monge patch

Tangent lines define an intrinsic metric expressed as *I* = *Edu*^2^ + 2*Fdudv* + *Gdv*^2^ where *E, F, G* are coefficients of the first fundamental form (Koenderink 1990). The second fundamental form defines tangent planes as change in surface shape of the normals with respect to the tangents (do Carmo 1976; Koenderink 1990) and are given as *II* = *Ldu*^2^ + 2*Mdudv* + *Ndv*^2^ (do Carmo 1976).

For a Monge patch on a surface, tangent vectors, **r** (*u,v*) = (**r**_*u*_, **r**_*v*_) = **r**_*uv*_, span the tangent plane and serve as a basis in a matrix of coefficients for the differential of the tangent vectors. For **r**_*uv*_ (*x*) = *u*_*r*_ **x**_*u*_ + *v*_*r*_ **x**_*v*_, **r**_*uv*_ (*y*) = *u*_*r*_ **y**_*u*_ + *v*_*r*_ **y**_*v*_, and **r**_*uv*_ (*z*) = *u*_*r*_ **z**_*u*_ + *v*_*r*_ **z**_*v*_, coefficients of the first fundamental form are *E* = **r**_*u*_ · **r**_*u*_, *F* = **r**_*u*_ · **r**_*v*_, *G* = **r**_*v*_ · **r**_*v*_, and ‖**r**_*u*_ × **r**_*v*_‖= (**r**_*u*_ · **r**_*u*_)(**r**_*v*_ · **r** _*v*_) – (**r**_*u*_· **r**_*v*_)^2^ = *EG* – *F*^2^. The normal to tangent vectors spanning a tangent plane is **n** (*u, v*) = (**n**_*u*_, **n**_*v*_) = **n**_*uv*_and the unit normal vector is 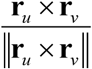. The coefficients of the second fundamental form are *L* = **r**_*uu*_ · **n** _*uv*_, *M* = **r**_*uv*_ · **n**_*uv*_, *N* = **r**_*vv*_ · **n**_*uv*_ (Koenderink 1990).

For the matrix of the differentials of the tangent vectors, the first and second fundamental forms have a determinant given as *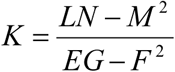*, and half the trace is *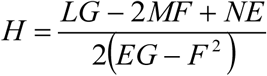* which are Gaussian and mean curvature, respectively (do Carmo 1976). Using *K*, shape can be determined as a general characterization of the surface (Koenderink 1990), and *H* is the overall shape of the surface as it is embedded in 3D space (do Carmo 1976).

#### 3D surface characterization via Gauss and Weingarten maps and the fundamental forms

The differentiable field of the unit normal vectors, *N*, on the surface, *S*, is a Gauss map of the tangent space of a point and is given as *N*: *S* → *S*^2^ for *S* → *R*^3^ and *S*^2^ = {(*x, y, z*) ∈ *R*^3^; |*x*^2^ + *y*^2^ += *z*^2^ = 1} (do Carmo 1976). The differential of a Gauss map is expressed as the shape operator or a Weingarten map and given as 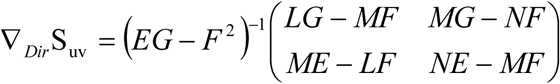 where 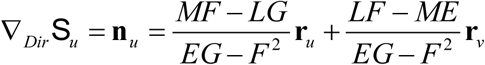 and 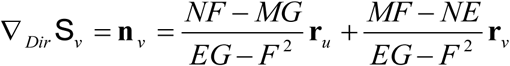. The shape operator determines all the tangent planes in the neighborhood of a point and is the change in surface normals with tangent vectors (Koenderink 1990).

#### Peaks, valleys and saddles of surface morphology and the Hessian

For a Monge patch with height function *z*,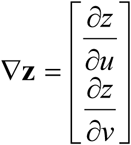 is the maximum height, and the magnitude is 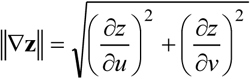 The Hessian is used to express the to the maximum slope at each point on the surface and is given as 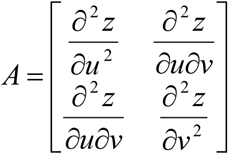. The product of the maximum height and maximum slope is expressed as 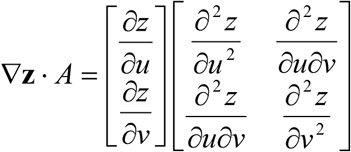. The characterization of extrema is accomplished by calculating eigenvalues of the Hessian, and the product of the eigenvalues is the Hessian determinant (Koenderink 1990). For each extremum on the surface of a Monge patch, the characteristic equation is **A h** = *φ* **h**, where *φ* is the eigenvalue of the Hessian expressed in matrix format as **A** and the column vector **h** represents ∇**z**. For the purpose of characterizing surface morphology using ∇**z** · *A*, eigenvalues are measures of the degree of concavity and curvature at a given *z*-value on the surface.

#### Smoothness as a characterization of surface morphology and the Laplacian

Characterizing the changes over the course of the surface—the ups and downs—is accomplished by the Laplacian (div (grad *z*)). For a patch, the Laplacian is 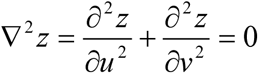 as the trace of the Hessian and a measure of the degree of average smoothness over the surface. The Laplacian is a harmonic function that is equal to the average of its nearby values on a surface. The Laplacian is also representative of steady-state or equilibrium conditions because the average smoothness is calculated at ∇^2^ *z* = 0 (Kaplan 2003).

To solve this initial value problem with a homogeneous boundary for a patch, separation of variables is used and expansion of ∇^2^ is a quadratic expressing each term as 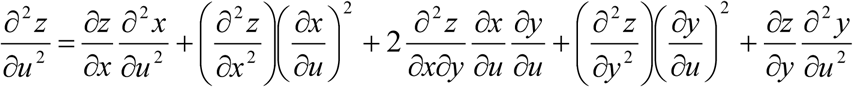 and 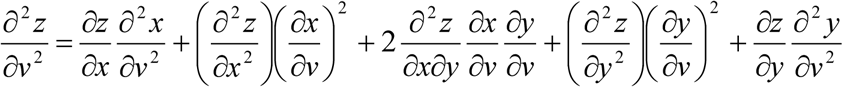. Discretization of the second partial derivatives with difference quotients (finite differences) determined by Taylor expansion of the Laplacian is given as 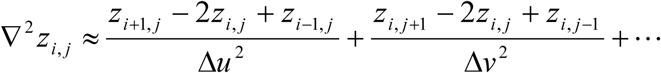 Each term is approximately equal to the eigenvectors of ∇^2^in *u* and *v* so that 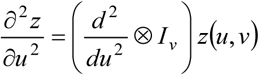 and 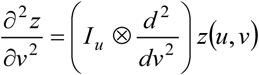. For ∇^2^*z*_*i,j*_, the eigenvalue problem is 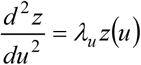 and 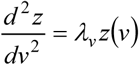 so that 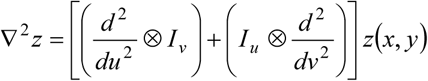 to determine degree of smoothness. The eigenvectors *eig* _*i, j*_ (*u, v*) = *eig* _*u*_ *⊗ eig* _*v*_ are 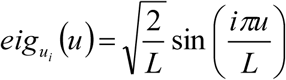 and 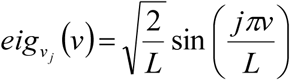 with Dirichlet boundary *∂*Ω = 0, and the eigenvalues are 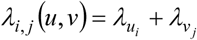, or more generally, *λ* = *λ*_*u*_ + *λ*_*v*_ (Weinberger 1965).

#### Point connections of 3D surface morphology and Christoffel symbols

The parameterization of a surface induces the connection between a 3D space and the embedding of 2D geometric structures in that 3D space. As an affine connection, a point is transported along a curve on the surface inducing a characterization of the geometry as tangent vectors of the surface in the neighborhood of that point in a tangent metric space. The derivative of tangent vectors of the surface induces a connection between tangent spaces that are nearby each other.

Christoffel symbols indicate where curvature is present in a locally flat neighborhood on a surface (Koenderink 1990). Values of Christoffel symbols that are non-zero are curvature indicators of degree of twisting, turning, contraction, or expansion of a surface locally about a given point; for vanishing or zero-valued Christoffel symbols, then flatness is defined in a local neighborhood of a given point (Misner 1973).

The metric tensor as a representation of the first fundamental form (do Carmo 1976) is a characteristic of differentiable inner products in tangent spaces, and because of this, Christoffel symbols can be expressed via the metric tensor components (Koop 1993; Kaplan 2003). Curvature is a characteristic of metric space as the metric tensor enables the calculation of distances (Sochi 2016) on space curves or surfaces embedded in 3D space. The metric tensor is *g*_*ij*_ = **e**_*i*_ · **e**_*j*_ where **e**_*i*_ and **e**_*j*_ are basis vectors in a coordinate system, and Christoffel symbols in basis *k* for these basis vectors are 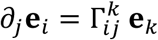 (Moore 2013). Surface basis vectors are contravariant space vectors or covariant surface vectors (Sochi 2016). A change in basis as a scalar product 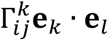 and *g*_*kl*_ = **e**_*k*_ · **e**_*l*_ produces 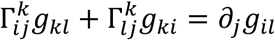 (Moore 2013). Basis vectors can vary from point to point on a curve or surface, and permutated indices are used to generate multiple Christoffel symbols in terms of the metric tensor (Kaplan 2003).

The addition of indices with respect to covariant and contravariant vectors undergoing differentiation will yield covariant and contravariant derivatives such as 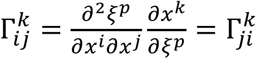 where *ξ*^*p*^ are standard coordinates from a fixed coordinated system, and the Jacobian is 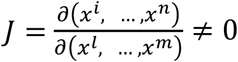 for the two sets of coordinates in bases *i* and *j* (Kaplan 2003) with *J*^*T*^*J* = *g*_*ij*_ as a contravariant component of the metric tensor. The Jacobian is related to Christoffel symbols as a generalized gradient (Misner 1973). While the Jacobian is undefined at zero (Kaplan 2003), Christoffel symbols at zero are indicators of flatness of the surface (Misner 1973).

The contravariant and covariant components of the metric tensor are expressed in terms of the standard coordinates as 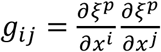 and 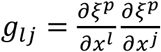 so that 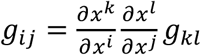 and 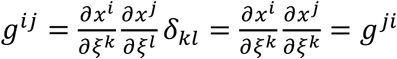, respectively, where *δ* is Kronecker delta (Sen and Powers 2012). Permutation of indices produces 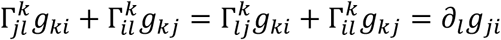 and 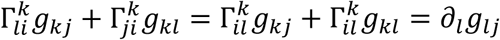 (Deserno 2004; Moore 2013). Collecting metric tensor components from the Christoffel symbols, and because 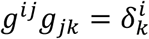 (Moore 2013), Christoffel symbols can be expressed in terms of the metric tensor components as 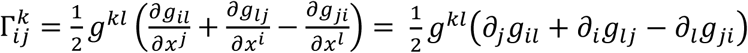 (Kaplan 2003; Moore 2013).

The first fundamental form coefficients and partial derivatives that are related to the metric tensor are used to express the connections between tangent spaces and can be summarized in matrix form as 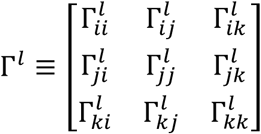 (Weisstein 2002). Christoffel symbols of the first kind via covariant surface basis vectors (Sochi 2016) are expressed in terms of the first fundamental form and its partial derivatives as 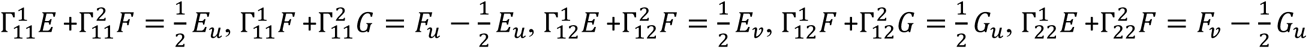 and 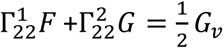 (do Carmo 1976; Sochi 2016). Christoffel symbols of the second kind via contravariant surface basis vectors are given as 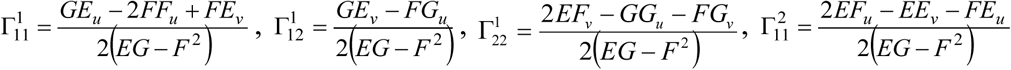, 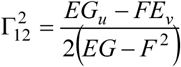 and 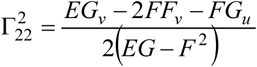 with 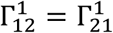 and 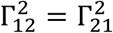 resulting from a torsion-free orientation of a space curve with respect to a Serret-Frenet frame at a given point on the surface (do Carmo 1976; Sochi 2016). The covariant surface metric tensor is 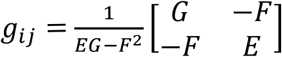, and the contravariant surface metric tensor is 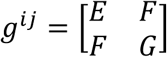 (Sochi 2016).

For a parametric surface, **r**(*u,v*)=(*x*(*u,v*),*y*(*u,v*),*z*(*u,v*)) 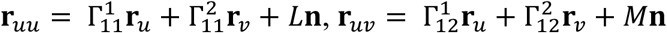 and 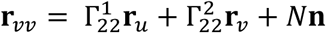 with (**r**_*uu*_)_*v*_ = (**r**_*uv*_)_*u*_ *and* (**r**_*vv*_)_*u*_ = (**r**_*uv*_)_*u*_ via Clairaut’s Theorem, indicating the relation between Christoffel symbols and the second fundamental form (Koenderink 1990). Additionally, Peterson-Codazzi-Mainardi equations relate Christoffel symbols to the second fundamental form as 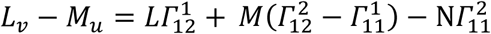 and 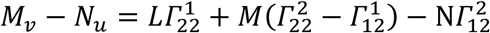 (Koenderink 1990).

Monge patches are local surfaces in a 3D metric space (Koenderink 1990). To calculate Christoffel symbols for a Monge surface patch via the first fundamental form, a change of variables via the total derivative is necessary because of the parameterization of the surface (Chase 2012). From elements of the Jacobian, the first fundamental form coefficients are 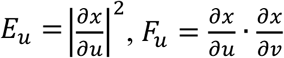, and 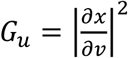 for parameter *u*, and 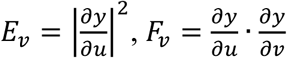 and 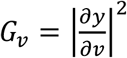 for parameter *v* (Kaplan 2003), and 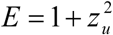 *F* = *z*_*u*_ *z*_*v*_, and 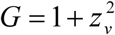 (Koenderink 1990). A Monge patch with height function *z* may be expressed in matrix form as 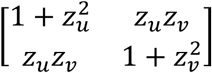, and 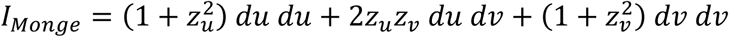 is the first fundamental form (Sochi 2016). For a Monge patch, the second fundamental form is 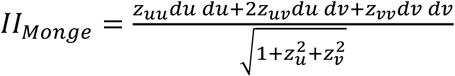 (Sochi 2016), and coefficients of the second fundamental form are expressed as 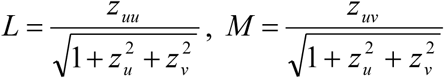, and 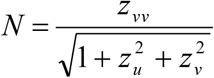 (Koenderink 1990). Christoffel symbols for a Monge patch may be calculated as 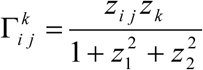 (Weisstein 2002) and represent the intrinsic geometry locally of curves and surfaces and their measurable characteristics such as the distance metric (Carmo 1976; Koenderink 1990) to connect surface features quantifiable on a pointwise basis.

Christoffel symbols of the second kind are the 2D surface connection coefficients of a 3D space (Sochi 2016), and the coefficients are a quantification of the degree of twisting, turning, contraction, or expansion of the surface. That is, the Christoffel symbols as connection coefficients serve three purposes: they are a quantification of how fast the components of a vector turn, twist, contract, or expand to keep constant against the turning, twisting, contraction, or expansion of the basis vectors; they are a quantification of parallel transport of basis vectors along the components of a vector; they are a quantification of the flatness of a surface (Misner 1973).

### Protocol for using centric diatom 3D surface models and their ensemble surface features in valve formation analysis

Parametric 3D surface models of external and forming valves of *Actinoptychus senarius, Arachnoidiscus ehrenbergii* and *Cyclotella meneghiniana* are used in analyses following Schmid and Volcani’s (1983) schema for centric diatom valve formation. For the modeling process, the final external valve model has *n*-horizontal slices of its surface removed in a stepwise fashion so that precursor steps are created to comprise the valve formation sequence. The ornamentation of the external valves is removed numerically at each step to illustrate the changes in valve formation and surface features. Models of each step for each taxon are devised using systems of parametric 3D equations. The number of steps chosen to create back-tracked slices of the valve formation in 3D is a function of the numerical changes in coefficients in systems of parametric 3D equations. The number of parametric 3D equations in each system representing a single diatom taxon is arbitrary because numerical solution can be obtained for any incremental, even infinitesimal, numerical change as the geometric change of a 3D surface. That is, the number of modeled steps potentially may be ad infinitum, even though perceptible changes are not evident.

Changes in ensemble surface features are used to develop a valve formation sequence as the morphogenetic model. Changes are measured as differences from one step to the next, starting with a base layer and sequentially accumulating ensemble surface features to complete the valve morphology. The base layer will have the smallest values for ensemble surface features so that, theoretically, the start would be a blank disk. The subsequent layers representing different degrees of sloping, peaks, valleys and saddles, smoothness, and flatness or curvature increases to model the acquisition of surface features to arrive at the final valve morphology. The whole surface is treated as a Monge patch when calculating ensemble surface measures.

Specifically, ensemble surface features are measured via first partial derivatives as elements of the Jacobian of a whole parametric 3D surface (variables *x, y* and *z* in parameters *u* and *v*), second partial derivatives as the elements of the *x*-, *y*- and *z*-Hessians of a Monge patch (where height is *z* with respect to planar patch *x*-*y*), the Laplacians as the sums of the diagonal elements of the *x*-, *y*- and *z*-Hessians, and Christoffel symbols as connection coefficients with contravariant indices *k* = 1, 2, *or* 3 and covariant indices *i, j* = 1, 2, 3. That is, Christoffel symbols of the second kind are calculated.

For each taxon, resultant quantified ensemble surface measures are analyzed for their contribution to each valve formation step and depicted in 100% stacked area plots to infer trends over time. The 100% stacked area charts show the contribution of each ensemble surface feature to each valve formation step. Each ensemble surface measure is analyzed to determine its contribution to ensemble surface features for *Actinoptychus senarius, Arachnoidiscus ehrenbergii*, and *Cyclotella meneghiniana* as well as the role each surface measure contributed in a comparison among valve formation taxon sequences. Total contribution of all Christoffel symbols to combined valve formation steps for all taxa was analyzed, and ensemble surface features for all taxa were analyzed with regard to ensemble surface measures as surface descriptors. A morphospace was devised to illustrate the relation between ensemble surface measures of the centric diatom taxa *Actinoptychus senarius, Arachnoidiscus ehrenbergii* and *Cyclotella meneghiniana* and changes during valve formation

## Results

Parametric 3D equations were used to construct models of *Actinoptychus senarius, Arachnoidiscus ehrenbergii*, and *Cyclotella meneghiniana* and their valve formation steps. Five valve formation steps for each taxon were modeled to depict noticeable valve face changes to fit into Schmid and Volcani’s (1983) three stage schema and are depicted in Figs. 2, 3 and 4. From single linkage cluster analysis using Hamming distance of non-zero values for ensemble measures, Schmid and Volcani’s (1983) stage 1 was covered by modeled valve formation steps 1 and 2, their stage 2 was covered by modeled valve formation steps 3 and 4, and their stage 3 was covered by the modeled final valve (Fig. 5). The number of modeled steps is arbitrary because clustering enables the binning of such steps into Schmid and Volcani’s tripartite schema. Five back-traced modeled morphologies were chosen only to enable perceptible change in valve formation steps. Ensemble surface features of the Jacobian, Hessian, Laplacian, and Christoffel symbols were calculated for all 3D surface models constructed from systems of parametric 3D equations. Christoffel symbols of the second kind are calculated (Hartle 2018) and are holonomic. For contravariant indices *k* = 1, 2, 3, covariant indices *i, j* are symmetric so that 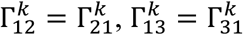, and 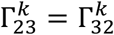.

**Fig. 2.**
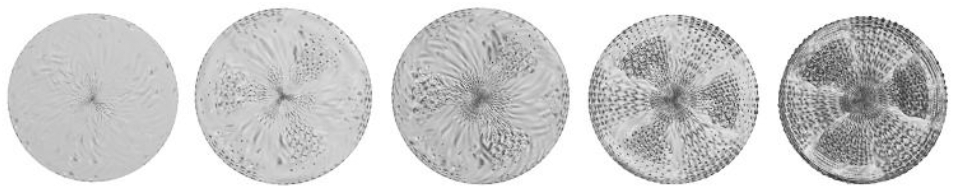
*Actinoptychus senarius* modelled valve formation sequence steps 1 through 5. From left to right, steps 1 and 2 represent stage 1, steps 3 and 4 represent stage 2, and step 5 as the finished valve represents stage 3 of Schmid and Volcani’s (1983) valve formation schema.

**Fig. 3.**
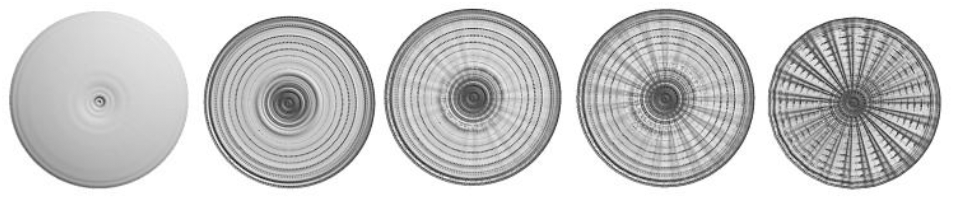
*Arachnoidiscus ehrenbergii* modelled valve formation sequence steps 1 through 5. From left to right, steps 1 and 2 represent stage 1, steps 3 and 4 represent stage 2, and step 5 as the finished valve represents stage 3 of Schmid and Volcani’s (1983) valve formation schema.

**Fig. 4.**
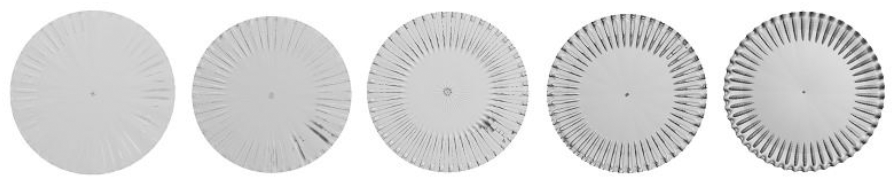
*Cyclotella meneghiniana* modelled valve formation sequence steps 1 through 5. From left to right, steps 1 and 2 represent stage 1, steps 3 and 4 represent stage 2, and step 5 as the finished valve represents stage 3 of Schmid and Volcani’s (1983) valve formation schema.

**Fig. 5.**
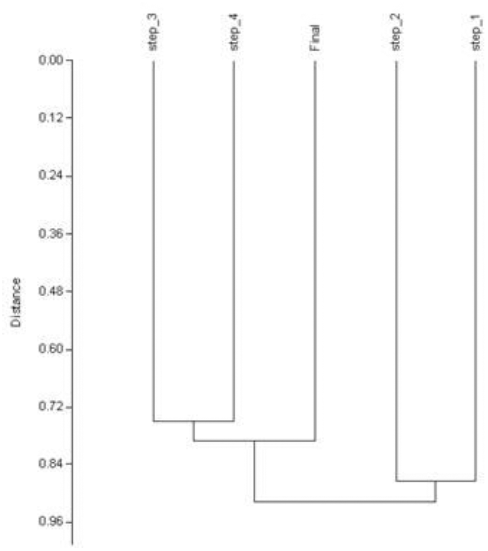
Single-linkage cluster analysis using Hamming distance for non-zero values of the Jacobian, Hessian, Laplacian, and Christoffel symbols.

For all models, the same *x*- and *y*-equations are used and are given as 16cos*u* cos *v* (1 + sin *u*) and 16 cos *u* sin *v* (1 + sin *u*), respectively. Only the *z*-equation differs for each taxon and are given in Table 1. Because the *x*- and *y*-equations were the same for all 3D models, each model can be treated as a Monge patch. As a result, the *z*-equation results are analyzed for the Jacobian in the *u*- and *v*-directions and the Hessian in the *u*-, *v*- and *uv*-directions. Christoffel symbols with contravariant index *k* = 3 are used for comparison to the Jacobian and Hessian *z*- equation values and the Laplacian. All Christoffel symbols are used in further analyses.

**Table 1.** Terms with parameters and coefficients of the *z*-equations for centric diatom taxa and valve formation steps. For all parameters, *u, ν* ∈ [0, 2*π*].

| Taxon | $z$-equation<sup>†</sup> |
| --- | --- |
| *Actinoptychus senarius* | $-\alpha u \cos 3u \cos 3v + \beta \sin 3u - \gamma \sin(6u)^5$ $-\zeta u \cos(5 - 4u) \sin(3 - 4v)^4 + \mu \sin 120u \sin 6v$ $-\varphi \cos(0.5u)^2 \cos(40u)^4 \cos 3v \sin 40u \sin(40v)^3$ $-\psi u \sin 40u \sin 160v$ |
| *Arachnoidiscus ehrenbergii* | $v \cos(0.5u^2) + \rho \cos(80u)^2 \sin(1.9u)^2 + \Xi \sin(1 + u)^3$ $+\tau \cos(10 + 74u^2) \sin(0.5 + 0.8u)^3 \sin(2 + 2.9u)^3$ $+\vartheta \cos u \sin(0.37u)^3 \sin(22v)^3$ |
| *Cyclotella meneghiniana* | $\cos 1.5u[\xi \cos 2u \sin 0.5u \sin 2.9u$ $+ 0.14 \cos u \sin(0.5u)^4 \sin(7v)^4]$ $+\eta \cos(3.5u)^6 \cos 25v \sin 1.8u^6 \sin 25v$ |

<sup>†</sup> Coefficients: $0.0008 \leq \alpha \leq 0.64$ , $0.01 \leq \beta \leq 0.1$ , $0.001 \leq \gamma \leq 1$ , $0.003 \leq \zeta \leq 0.09$ , $0.0025 \leq \mu \leq 0.125$ , $0.2 \leq \varphi \leq 10$ , $0.0025 \leq \psi \leq 0.025$ , $0.004 \leq \vartheta \leq 0.7$ , $0.05 \leq v \leq 0.75$ , $0.025 \leq \rho \leq 0.5$ , $0.051 \leq \tau \leq 2.55$ , $2 \leq \Xi \leq 3$ , $0.01 \leq \xi \leq 0.5$ , $0.18 \leq \eta \leq 3$ .

A combination of all the ensemble measures of the Jacobian, Hessian, Laplacian, and Christoffel symbols produced the depiction of change from the start to each subsequent step of modeled valve formation with regard to the *z*-term and Christoffel symbols for contravariant index *k* = 3. The Jacobian in *u*- and *v*-parameters and the Hessian in the *u*-parameter were positive influences with increasingly positive contribution from start to final valve for *Actinoptychus senarius* (Fig. 6). Negative influences were evident for the Hessian *v*- and *uv*-parameters, the Laplacian as well as Christoffel symbols 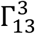 and 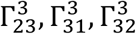. Largest negative input was the Hessian *v*- and *uv*-parameters and Laplacian for step 4 and the final valve and Christoffel symbols 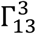 and 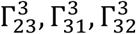 for steps 1 and 3 for *Actinoptychus senarius* (Fig. 6).

**Fig. 6.**
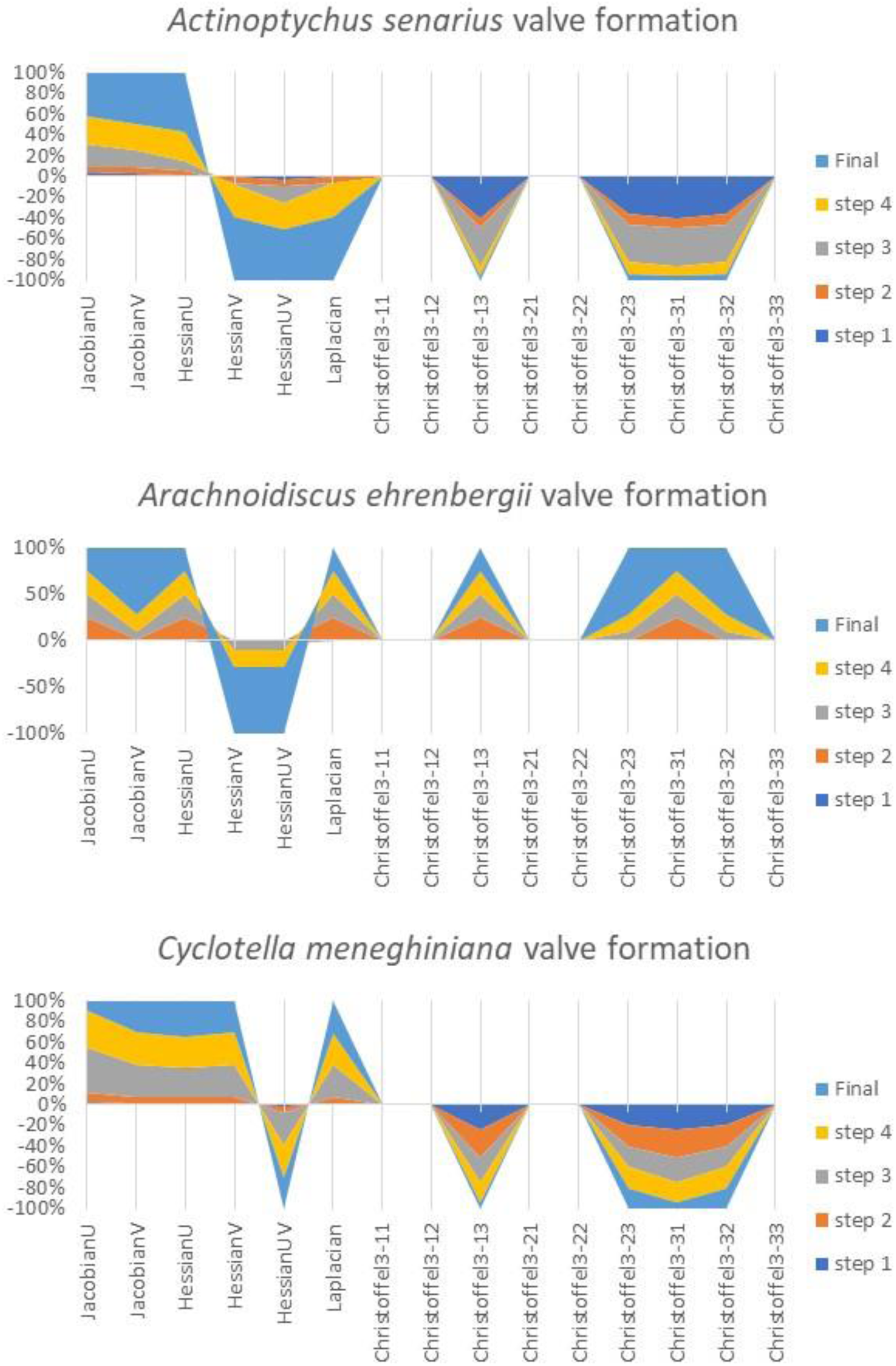
Contribution percentage of ensemble surface measures Jacobian, Hessian, Laplacian, and Christoffel symbols to each valve formation step.

For the Jacobian in the *u*-parameter and Hessian in all parameters, the scenario for *Arachnoidiscus ehrenbergii* was similar to that for *Actinoptychus senarius* (Fig. 6). The Jacobian in the *v*-parameter had a near zero value for steps 1 thorough 4. Only the final valve had a positive value. The Laplacian and Christoffel symbols 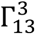 and 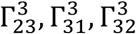 were positive influences, contributing almost equally to steps 2 through the final valve for *Arachnoidiscus ehrenbergii*, with Christoffel symbols 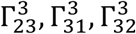 contributing a larger portion to the final valve (Fig. 6).

*Cyclotella meneghiniana* had positive influence from the Jacobian and Hessian in the *u*- and *v*-parameters and the Laplacian (Fig. 6). The Hessian in the *uv*-parameter and Christoffel symbols 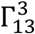 and 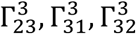 were negative influences in almost equal contribution for steps 1 through the final valve, with the exception of almost no contribution by 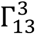 for the final valve of *Cyclotella meneghiniana* (Fig. 6).

Modeled taxa were analyzed by their valve features using each ensemble surface measure. The ensemble surface measures that were most indicative of peak height changes in *Actinoptychus senarius* valve features were the Hessian in the *v*- and *uv*-directions from step 2 through the final valve, reflecting the changes in alternating undulations of the six valve surface sectors radiating outward from the center (Figs. 7d, f). Only a slight uptick in value from step 3 to step 5 for the Jacobian in the *v*-direction was evident, indicating the change in sloping of undulations from sector to sector on the valve surface (Fig.7b). All other ensemble surface measures including the Jacobian and Hessian in the *u*-direction, Laplacian, and Christoffel symbols for contravariant index *k* = 3, indices *i* = 1, 2 and *j* = 3 were near zero, indicating an overall regular valve pattern of the smaller surface features in contrast to the larger valve features of the alternating undulations of the six sectors (Figs.7a, c, e, g and h).

**Fig. 7.**
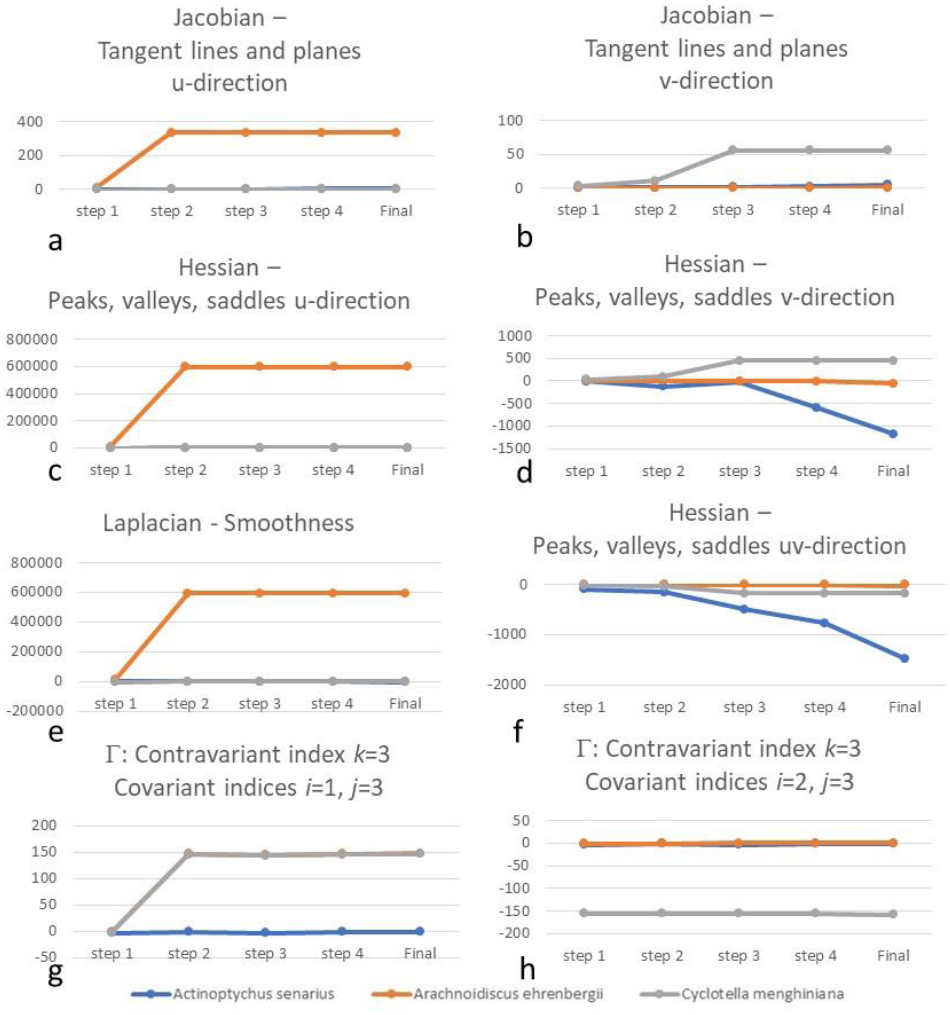
Jacobian, Hessian and Laplacian from *z*-terms and Christoffel symbols 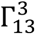 and 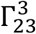 for modelled valve formation steps 1 through the final valve for *Actinoptychus senarius, Arachnoidiscus ehrenbergii* and *Cyclotella meneghiniana*.

*Arachnoidiscus ehrenbergii* exhibited large surface changes from valve formation step 1 to step 2 as sloping and peakedness as the Jacobian and Hessian in the *u*-direction (Figs. 7-a, c). Regular undulations from the ribs to the concentric circles below the ribs is recovered by the Laplacian as “harmonics” (Fig. 7-e). While curvature changes in the ribs differing from the underlying concentric rings on the valve face are indicated by Christoffel symbols for contravariant index *k* = 3, indices *i* = 1 and *J* = 3 (Fig. 7g), the Jacobian in the *v*-direction, Hessian in the *v*- and *uv*-directions, and Christoffel symbols for contravariant index *k* = 3, covariant indices *i* = 2 and *j*= 3 are near zero, indicating transitional or saddle areas of the valve surface that are flat (Figs. 7d, f, and h). *Arachnoidiscus ehrenbergii* has a fairly even, regular patterned surface as measured by ensemble surface features.

The slightly undulating surface of *Cyclotella meneghiniana* was reflected in the Jacobian in the *v*-direction, increasingly from step 1 to step 3 during valve formation (Fig. 7b), in contrast to the *u*-direction (Fig. 7a). The same pattern held for the Hessian in the *v*-direction with positive values indicating valleys (Fig. 7d) as well as Christoffel symbols for contravariant index *k* = 3, covariant indices *i* = 1 and = 3 indicating greater curvature areas (Fig. 7g). Sharp changes in peak height as an outer ring of “pleats” or plications at the valve margin of *Cyclotella meneghiniana* are recovered via these ensemble surface measures. A slight depression in the general valve surface was indicated by the Hessian in the *uv*-direction (Fig. 7f) representing the central area, and a transitional or saddle area between peaked and depressed areas along the marginal plications was indicated by the Hessian in the *u*-direction (Fig. 7c). Valve surface smoothness is reflected in the very lightly radially streaked central area via the Laplacian (Fig. 7e), while flatness of the overall valve face shape was indicated by Christoffel symbols for contravariant index *k* = 3, covariant indices *i* = 2 and = 3 (Fig. 7h).

The *u*- and *v*-parameters of the *z*-term in the Jacobian and Hessian and sum of the *z*-terms in the Laplacian contributed differing amounts to the outcome of valve formation for each taxon. For the Jacobian, all steps of *Cyclotella meneghiniana* were influenced by the *v*-parameter, and all steps of *Arachnoidiscus ehrenbergii* were influenced by the *u*-parameter as was the case for *Actinoptychus senarius*, but to a lesser degree (Fig. 8, top). For the Hessian, the *uv*-parameter had a larger influence than the *v*-parameter on all steps of *Cyclotella meneghiniana* and *Actinoptychus senarius*. For *Cyclotella meneghiniana*, the influence was positive, while for *Actinoptychus senarius* the influence was negative (Fig. 8, middle). *Arachnoidiscus ehrenbergii* was influenced by the Hessian *z*-term in the *u*-parameter ((Fig. 8, middle). The Laplacian was a negative influence for *Actinoptycus senarius* and a positive influence for *Cyclotella meneghiniana* and *Arachnoidiscus ehrenbergii* (Fig. 8, bottom).

**Fig. 8.**
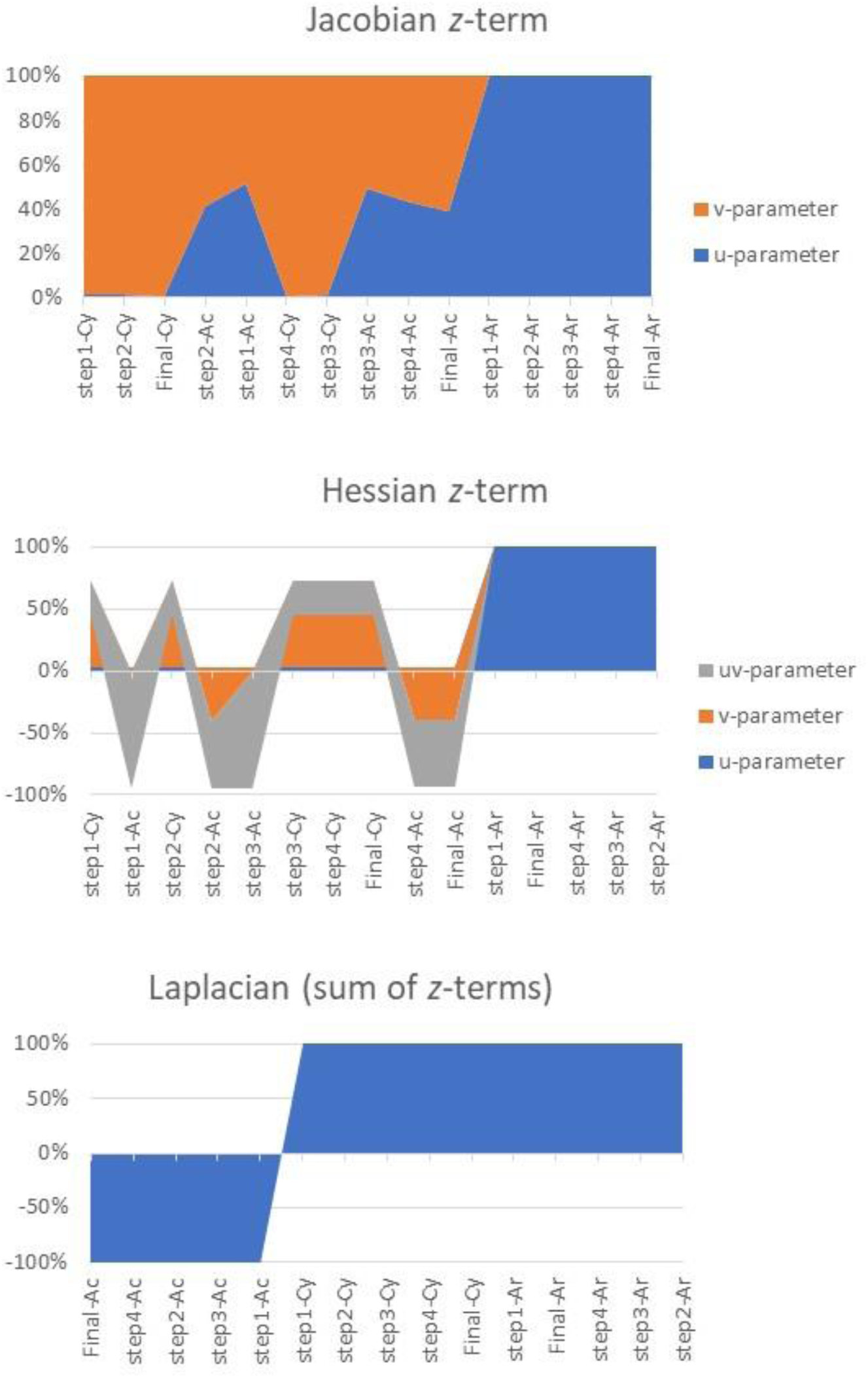
Contribution of the *u,v*-parameters of the *z*-term of the Jacobian, Hessian and Laplacian to each step of valve formation for *Actinoptychus senarius, Arachnoidiscus ehrenbergii* and *Cyclotella meneghiniana* expressed as a percentage.

All Christoffel symbols were analyzed to determine degree of flatness and to determine their contribution to valve characteristics at each step of valve formation on the valve surface. The point at which the Christoffel symbols vanish are flat areas in a torsion-free or holonomic frame. The direction of change in a tangent space is symmetric in the covariant indices when Christoffel symbols equal zero. The non-zero values indicate distortion resulting from curvature as the connections on the valve surface change from point to point.

For contravariant index *k* = 1, steps 2, 3, 4, and the final valve were represented as a large percentage of negative values of 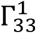, then transitioning to step 1 as a narrower percentage of positive values for step 1 of *Arachnoidiscus ehrenbergii* (Fig. 9-top). Step 1 through final valve for *Actinoptychus senarius* and *Cyclotella meneghiniana* were represented as a large percentage of positive values of 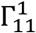 (Fig. 9-top). All other values of Christoffel symbols for contravariant index *k* = 1 were equal to or near zero.

**Fig. 9.**
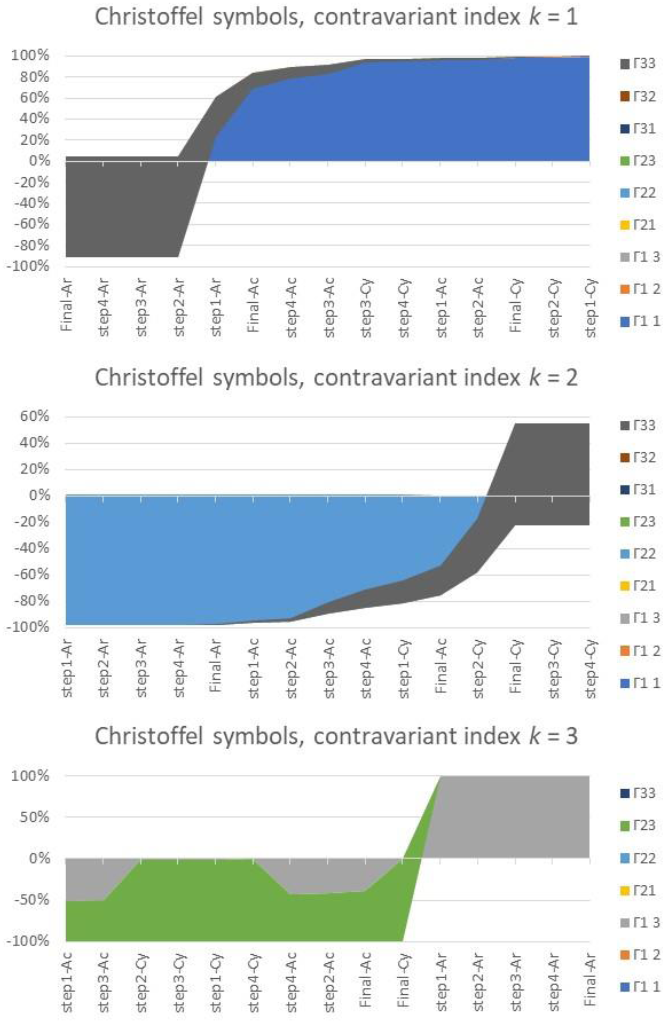
All Christoffel symbols for the three contravariant indices: top, *k* = 1; middle, *k* = 2; bottom, *k* = 3. Shaded areas indicate percentage that a given Christoffel symbol contributed to a valve formation step for each taxon. Because of symmetry, Christoffel symbols with contravariant index *k* = 3, covariant indices *i* = 3, *j* = 1 and *i* = 3, *j* = 2 are not reported.

For contravariant index *k* = 2, step 1 through the final valve for *Arachnoidiscus ehrenbergii* had a large percentage of negative values for 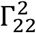 (Fig. 9-middle). For step 1 through the final valve of *Actinoptychus senarius* and steps 1 and 2 for *Cyclotella meneghiniana*, a larger percentage of negative values of 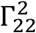 was indicated over a smaller negative percentage for 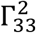 (Fig. 9-middle). For *Cyclotella meneghiniana* steps 3, 4 and the final valve, only 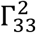 had a larger percentage of positive versus a lesser percentage of negative values (Fig. 9-middle). All other values of Christoffel symbols for contravariant index *k* = 2 were equal to or near zero.

The general order of the taxa was switched for contravariant index *k* = 3 Christoffel symbols. All steps for *Actinoptychus senarius* and *Cyclotella meneghiniana* were large percentages of negative values for 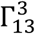 and for 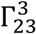, where the latter indicated the bulk of the negative values (Fig. 9-botton). By contrast, all steps of *Arachnoidiscus ehrenbergii* were a large percentage of positive values for 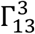 and a smaller percentage of positive values for for 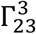 (Fig. 9-bottom).

All Christoffel symbols for all three taxa were plotted for the sequence of steps in valve formation. Each step is covered by all contravariant indices *k* = 1, 2, 3. Of all the Christoffel symbols, those that were non-zero and had contributed to non-flat areas of the valve face were 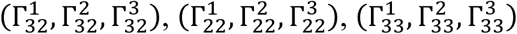, and 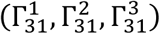. The contribution of Christoffel symbols with covariant indices *i* = 3, *j* = 2 was a constant negative value for steps 1 through the final valve. Covariant indices *i* = 2, *j* = 2 had a similar contribution. On the positive side, covariant indices *i* = 3, *j* = 1 contributed to valve formation steps 2 through the final valve as a constant. Only step 1 had no contribution from Christoffel symbols with covariant indices *i* = 3, *j* = 1. For Christoffel symbols with covariant indices *i* = 3, *j* = 3, their contribution was exponential from step 1 to step 3 then levelled off as a maximum for steps 3, 4 and the final valve (Fig. 10).

**Fig. 10.**
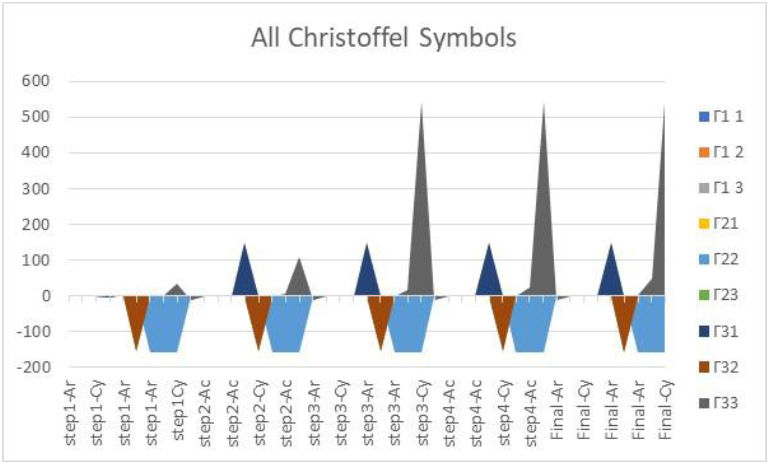
Plot of all Christoffel symbols shows change in contribution for 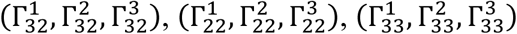, and 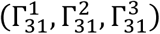 per valve formation step.

To aggregate ensemble surface measures and their contribution to valve formation steps for all taxa combined, a plot was devised to show the percentage of that contribution in terms of the descriptors flatness, peaked, sloped, and smoothed (Fig. 11). Contraction described step 1, almost exclusively. Contraction to a lesser degree described step 2 with smaller contributions from almost flat to expansion, peaked and sloped features, and a slight contribution of dipped features. Valve formation step 3 included contraction, almost flat, expansion, dipped, peaked, and sloped descriptors along with the addition of the concave smoothness descriptor. Step 4 had elements of the same contributing descriptors with the addition of convex smoothness. The final valve was similar to step 4 but had the least contribution from the contractor descriptor compared to the previous four steps in the valve formation sequence (Fig. 11).

**Fig. 11.**
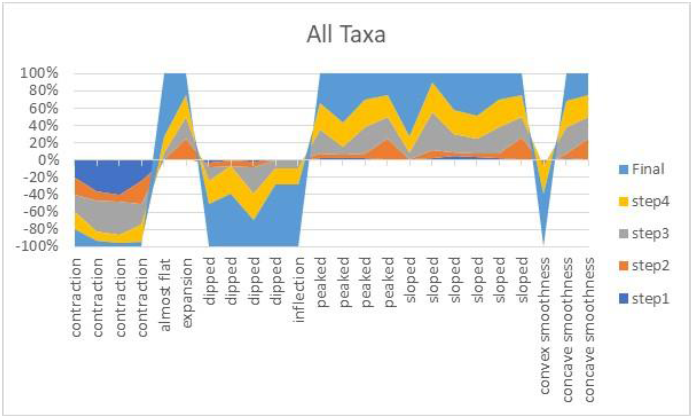
Ensemble surface measures as indicators of ensemble surface features for each valve formation step.

A 3D morphospace was devised for *Actinoptychus senarius, Arachnoidiscus ehrenbergii* and *Cyclotella meneghiniana* using ensemble surface measures of valve formation (Fig. 12). Peakedness and smoothness were combined on one axis because the trace of the Hessian is the Laplacian. Slopeness and flatness were combined on another axis because differentiation of the Jacobian is a coordinate transformation of non-vanishing Christoffel symbols (Bertschinger 1999).

**Fig. 12.**
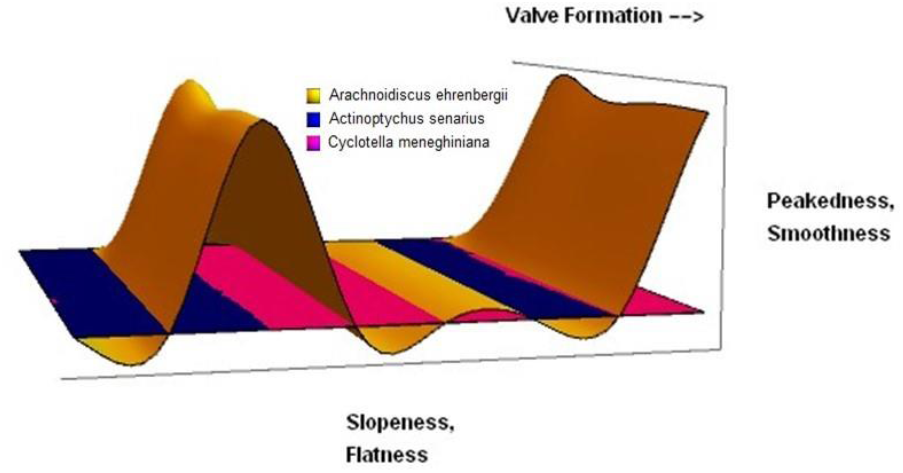
A 3D morphospace of ensemble surface measures of valve formation for *Actinoptychus senarius, Arachnoidiscus ehrenbergii* and *Cyclotella meneghiniana*.

The morphospace shows a “cut-away” space to facilitate interpreting taxon ensemble surface feature results. The most pronounced changed in ensemble surface features occurred for *Arachnoidiscus ehrenbergii*. Emanating from its center, *Arachnoidiscus ehrenbergii* has alternating ribs and dips as its valve surface, exhibiting peaks and valleys. The ribs exhibit smoothness, while the branching or reticulated pattern beneath the ribs exhibits peakedness. The valve surface pattern exhibits a regular fluctuation in a harmonic fashion (Fig. 12). During valve formation, *Arachnoidiscus ehrenbergii* proceeds with more changes in peakedness and smoothness and less so concerning slopeness and flatness (Fig. 12).

Because the scale at which *Arachnoidiscus ehrenbergii* ensemble surface features differed was so great, a second morphospace “cut-away” was devised to enable the visualization of details for *Actinoptychus senarius* and *Cyclotella meneghiniana* (Fig. 13). A marked change in peakedness and smoothness more so slopeness and flatness was evident for *Actinoptychus senarius*. A sharp dip proceeding from the first step to final valve in the valve formation sequence suggested the formation of the alternating undulations of the sectors on the valve surface (Fig. 13).

**Fig. 13.**
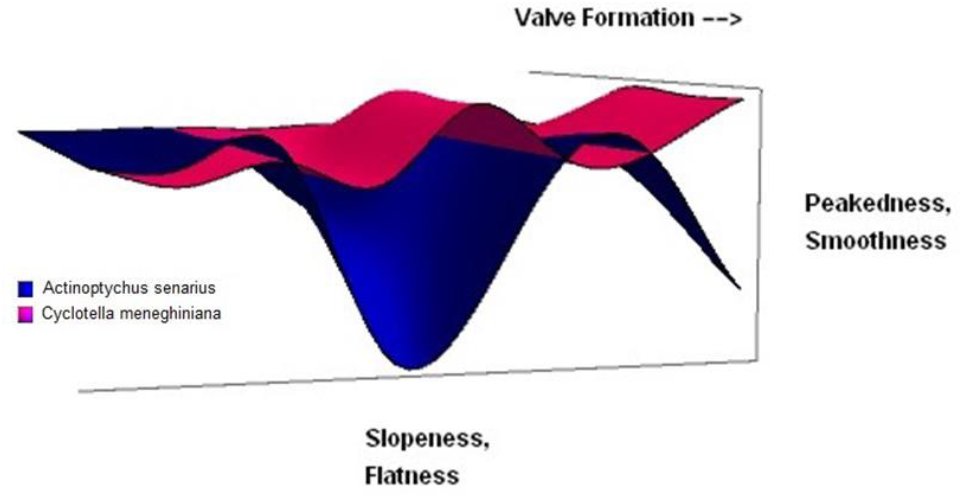
A 3D morphospace of ensemble surface measures of valve formation for *Actinoptychus senarius* and *Cyclotella meneghiniana*.

For *Cyclotella meneghiniana*, the changes in peaks and valleys as well as slopeness and flatness varied in a shallow harmonic fashion (Fig. 13). The shallow depth of the change in peakedness and smoothness suggested the finer undulation of the valve marginal plications in contrast to the regular harmonic undulating changes in the valve surface of *Arachnoidiscus ehrenbergii* or the large undulating sectors of *Actinoptychus senarius*. The slightly larger peak with respect to slopeness and flatness suggested the presence of the central area of *Cyclotella meneghiniana*. For the valve formation sequence, progression from more slopeness and flatness to more peakedness and smoothness indicated the final valve marginal surface of *Cyclotella meneghiniana* (Fig. 13).

In the morphospace, the areas where smoothness equals flatness are inflection points that may be saddle points as well. That is, at the intersection of the areas for *Actinoptychus senarius, Arachnoidiscus ehrenbergii* and *Cyclotella meneghiniana*, the value equals or is very close to zero (Fig. 13).

## Discussion

Diatom morphogenesis is a compilation of silica deposition, valve formation, mitotic growth, and other cytological, genetic and epigenetic processes (e.g., Schmid and Volcani 1983). Silica deposition during morphogenesis is not uniform (Vartanian et al. 2009). Overall, valve surface pattern is maintained and reproducible, but diatom growth concerning silica deposition and the geometry of the surface occurs within finite spatial boundaries (Vartanian et al. 2009). Regarding phenotypic plasticity as an influence on diatom valve patterns along with processes governing valve formation cytologically, 3D surface models are useful in studying potential modes of growth in morphogenesis.

Centric diatom valve morphogenesis was explored using 3D surface models and measurements of boundaryless, variable morphological characters combined as ensemble surface features. Studies were conducted with regard to stepwise ensemble surface changes as a sequence of events during valve formation. Each of the three general stages of valve morphogenesis as defined by Schmid and Volcani (1983) were used as guidelines for the changes in ensemble surface features as slices in the height distribution of silica deposition. At the first stage of horizontal silica deposition and the presence of an annulus and branching pattern of silica strands with radial rows, this basal silica layer is the smoothest and flattest, having smaller values of slopes, peakedness and saddles. At the second stage of vertical silica deposition, areolae walls increase in height that is reflected in larger values of peakedness and saddles. At the third stage of increasing horizontal and vertical silica deposition, peakedness and saddles become more differentiated as the size and spacing of such structures may become exaggerated with intervening areas of smoothness and flatness, as silica deposition extends to the valve margin.

At each point on the valve surface, ensemble surface features may have positive, negative and zero values that are indicators of the degree of sloping, peakedness and saddles, smoothness, or flatness (Table 2). Positive values for elements of the Jacobian indicate increasing sloping or steepness of the surface, while negative values indicate decreasing sloping. Positive values for elements of the *z*-Hessian are peaks, negative values are valleys, and zero values are saddles. The transition from positive to negative values are indicators of changes in smoothness as harmonics, as smoothness is when the Laplacian is zero. Positive or negative Christoffel symbols are indicators of curvature, while flatness is recorded in zero values.

**Table 2.**
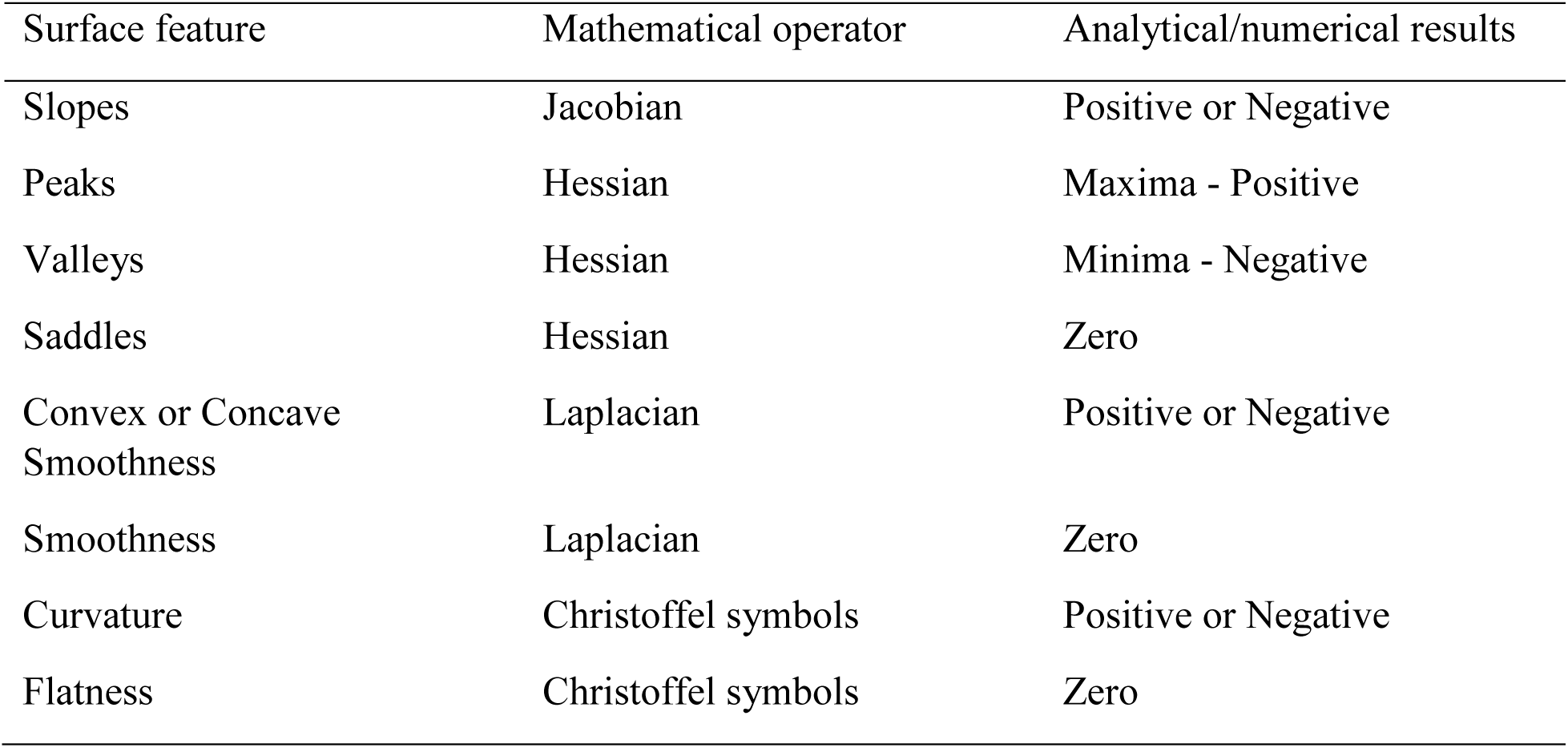
Ensemble surface features, mathematical operator and numerical results.

Ensemble surface measures from parametric 3D models were readily compiled and corresponded to Schmid and Volcani’s (1983) schema of diatom valve morphogenesis. Quantification of diatom surface morphology using such measures enabled the study of valve morphogenesis as well as showed the efficacy of using 3D surface models in valve formation studies. Such models can be used to extend the study of valve formation to include the initial cell and auxospore of the diatom life cycle. Systems of parametric 3D equations can be devised to create 3D surface models when initial cell and auxospore surface and shape morphology is known and documented.

### Ensemble surface features and physical characteristics of valve morphogenesis

Ensemble surface features may be present indicating rotational, reflective, dihedral, or conformal symmetry. The same features may indicate degrees of complexity with regard to degree of sloping, peakedness, smoothness, or flatness. Increasing sloping and peakedness measured as the Jacobian and Hessians, respectively, means increasing complexity in contrast to increasing smoothness or flatness as measured from the Laplacian and Christoffel symbols, respectively. Symmetry and complexity assessments at each stage in valve formation may be used as composite descriptors of the changes in development, and such changes may be related to the changes occurring at the subcellular level with regard to organelles and their movement during mitosis and the size diminution cycle for many diatoms (Pappas et al. 2019). Ensemble surface features may indicate whether circular centric diatoms can become asymmetric at a given stage of growth and valve formation despite being formed from an annulus. From stepwise valve formation via composite ensemble surface features, a general ranking of most to least symmetric taxa are *Arachnoidiscus* to *Actinoptychus* to *Cyclotella*. A composite of ensemble surface features indicates that the highest to lowest degree of complexity in diatom valve surface formation patterns is *Arachnoidiscus* to *Actinoptychus* to *Cyclotella*.

Valve formation is directional with respect to valve characters and pattern, but at a local level. Initially, branching patterns may exhibit waviness, but spatial constraints on silica deposition occur to induce regularly-spaced surface features, such as areolae, as wave front expansion from annulus to valve margin occurs in centric diatoms (Vartanian et al. 2009). Expansion of a wave front with respect to the SDV may induce fractal-like structures at the valve margin with respect to growth (Vartanian et al. 2009).

Such growth may be indicative of the first stage of valve formation. At the second stage of valve formation, silica fills in the valve face producing another layer of patterning which occurs behind the initial wave front expansion. One possibility is that vertical silica deposition might occur stochastically, resulting in changing sizes and shapes of pores and rimoportulae in a random fashion over the valve face and in contrast to horizontal silica deposition that occurs chaotically resulting in fractal-like structures. At the third stage of valve formation, a combination of stochastic and directed processes might occur as cribra and cribella form within the constraints of the areolae, but silica filling into gaps in the valve face structures may do so in a random fashion. This represents growth where cell division and symmetry occur as a propagation of wave fronts (Nechaev 2017).

### Factors affecting valve formation

Size and geometry of a morphological character may be affected by enzymatic inhibitors in contrast to the spatial distribution of those characters on the valve surface (Vartanian et al. 2009). At the first stage of valve formation, constraints on the valve surface may be laid down with regard to transapical costae and at the second stage of valve formation with the formation of cross costae in some centric diatom taxa (Vartanian et al. 2009). Sequential deposition with respect to silica height on the valve surface records pattern formation over time and may be affected by inhibitors. Low concentrations of inhibitor may induce change in silica deposition and a change in development of morphological characters on the valve. High concentration of inhibitors may induce aberrant valves (Vartanian et al. 2009).

Development of a morphological character is affected by inhibitors in contrast to the spatial relation among morphological character on the valve surface of a diatom (Vartanian et al. 2009). Fluctuations in processes at the molecular level may induce slight differences in silica deposition on different parts of the valve surface. Physically, exponential cell growth may produce excess silica material generation to induce buckling of the domain boundary (Nechaev 2017) or buckling on the surface. Structural anomalies such as areolar row disordering or distorted valve face shape may be induced via microtubule inhibitors because microtubule dissolution occurs or chromosome separation does not (Bedoshvilli et al. 2018). In either case, buckled valve structures may result as valve morphogenesis occurs.

Because valve morphogenesis commences at the microtubule center, and subsequent development of large valve structures (e.g., Pickett-Heaps 1998; Van de Meene and Pickett-Heaps 2002, 2004; Tesson and Hildebrand, 2010a, b) occurs in conjunction with microtubule activity, morphogenetic stages that are interrupted or changed by microtubule inhibitors may produce abnormal valve structures or valve shape (Bedoshvilli et al. 2018) as buckled surface or marginal valve margins. Such differences may be modeled using ensemble surface features with the potential of matching numerical results with the associated positioning of microtubules (e.g., Cohn et al. 1989) and possible valve surface structural or marginal buckling. The appropriate combination of such features has yet to be discerned and how such combinations might be associated to particular steps in the silica deposition process, and how this represents valve formation at given times during this process remains to be studied.

### Diatom growth patterns—buckling and wave fronts

Changes in early growth during diatom morphogenesis may resemble the propagation of an optical ray. Changes in the trajectory of the ray may resemble changes in the growth path during valve formation of diatom morphogenesis. For a homogeneous medium, that path may be linear and constant. Otherwise, the path may be non-linear but follow an exponential or logarithmic path. Rays normal to the surface are wave fronts (Ries and Muschaweck 2002) and may represent buckling patterns on diatom surfaces. A buckled surface is a patterned surface (Chen et al. 1998; Lin et al. 2000).

A wave front or buckling may be interpreted to occur on the surface of *Cyclotella meneghiniana* with regard to the changes from the central area to the margin of the entire valve. Another aspect of a wave front property may be construed to be in the pleating or plications near the valve margin with regard to the areolae structure in a ring. The undulating sectors of *Actinoptychus senarius* or the height changes between ribs and underlying structural branching pattern of *Arachnoidiscus ehrenbergii* may be representational of buckling that has wave front properties.

The Jacobian and Hessian elements of first and second partial derivatives, respectively, record the indications of buckling on the surface of a diatom valve. By contrast, the Laplacian and Christoffel symbols record the smoothness and flatness, respectively, of the diatom valve surface which indicates lack of buckling. As a physical phenomenon, buckling is non-uniform compression given as differential growth (Nechaev 2017). For isometric planar growth, buckling does not exist. That is, early growth from a conical surface above a disk corresponding to isometry is a constant growing surface where growth is low and the surface is flat, and buckling does not occur (Nechaev 2017). Ensemble surface features and the composite change in these features across a surface may be a variant of wave front propagation. The Laplacian has solutions that are representative of harmonics, and the combinations of first and second partial derivatives are extractable as variants of wave front solutions.

For an exponentially growing surface embedded in a plane, the Jacobian of a conformal mapping is dependent on angular symmetry and is recoverable in terms of hyperbolic metrics of the surface. This growth is a type of motion in which energy is propagated from one stage to another along a continuum from nascent cell to mature organism. Buckling as a wave front is a natural phenomenon of cell growth at different scales (Nechaev 2017).

Buckling on the surface may be determined by the metric tensor. Geodesics defining paths on the surface are parameterized and determine the Christoffel symbols. Geodesics along a surface are associated with the motion of silica deposition at regular intervals producing a characteristic valve pattern. Movement of silica with regard to a valve surface may be indicative of cell division and symmetry changes as a propagation of wave fronts. Surface height above the domain is characterizable by such wave front movement (Nechaev 2017). Degrees and kinds of surface buckling may be indicators of different rigidities of surface features (Gordon and Tiffany 2011).

At which junctures buckling occurs during diatom valve morphogenesis has yet to be determined (Gordon et al. 2009). One hypothesis is that buckling occurs when frustule structures are thin (Gordon and Tiffany 2011). Another hypothesis is that via wave front analysis, buckling may happen at any time during valve formation as a natural phenomenon that is genetically, environmentally, or epistatically controlled. This may induce plasticity in the phenotype. Epistasis may be instrumental in buckling as a more complicated, unpredictable, chaotic phenomenon. Ensemble surface measures may be used to characterize phenotypic buckling during morphogenesis because a 3D surface is a proxy for the phenotype (Pappas and Miller 2013).

At some critical values, the Jacobian may become negative and the resultant growth may be squeezed in rather than buckling out. Squeezing in may be a regulation of the rigidity or stiffness of the surface (Nechaev 2017) as in buckling, which is constrained by surface features that are ridged or stiff (Gordon and Tiffany 2011). Growth may be geometrically hyperbolic in which there is zero Gaussian curvature (Nechaev 2017). At constant negative curvature, the surface bending may be similar to a pseudosphere. Cascades of pseudospheres may resemble peaks and saddles, exhibiting buckling in this way. High positive values of the Hessian may measure buckling with respect to silica height off the valve surface, while the high positive values for the Laplacian and zero values for Christoffel symbols may be used to measure lack of buckling, or possibly, transitions between buckles that are not saddles on valve surfaces with few silica elevations as morphological characters. Locally on a valve surface, negative values of Hessians, Laplacians and Christoffel symbols may be indicators of squeezing out rather than buckling in. With the potential for a hierarchical, nested or cascading structure in terms of the relation among ensemble surface features, repetition of such features may be present at multiple scales.

Early in growth, there are buckling instabilities on the circumference as negative values. At a critical point on the surface, buckling proliferates in the direction of growth where peaks and valleys multiply. A hierarchy in peak size occurs related to degree of buckling. At later stages in growth, buckling instabilities subside as the limits of growth occur. At the circumference, a self-similar buckling profile may emerge (Nechaev 2017).

### Valve formation, ensemble surface features and self-similarity

Self-similarity may be evident during valve formation, but such a determination requires special testing. One possible test may be to detect the smallest changes among pairwise smallest *z*-Hessian elements and using ultrametricity (Nechaev 2017), find the smallest possible scale at which a given feature is measurable, and find the next larger scale at which that feature is manifested which may be present on the same valve face or on a valve face generated during valve formation. If a given feature is found at different scales, then there is self-similarity on the diatom valve surface at times when growth or silica deposition occurs. Self-similarity may be at work during growth at different scales and may be evident and measurable as scale symmetry. The limits to growth and the MacDonald-Pfitzer rule (MacDonald 1869; Pfitzer 1869; 1871) may determine the threshold at which self-similarity may or may not be evident.

### Diatom morphogenesis: cytoplasmic inheritance and phenotypic plasticity

Cytoplasmic inheritance is non-genetic inheritance in which there is a division of cell states between cell lineages (Shirokawa and Shimada 2016). Structure of the parental cell can be expressed by the daughter cell through direct descendance or there could be a pre-existing cell membrane that functions as a template for development of the daughter cell. Diatoms reproduce via daughter cells forming within the SDV that expands within the parent cell, decreasing the size of the offspring. Daughter cell microscale surface structure patterns (i.e., morphological characters such as central fultoportulae, striae, central area, and cell diameter) correlate with parental organelles (Shirokawa and Shimada 2016).

Cell structure traits are quantitatively different from structural traits inherited by genetic or non-genetic factors. In diatoms, cell structure variation results from the new formation of daughter cells within the parent as incomplete inheritance (Shirokawa and Shimada 2016). Microscale structural patterns on the daughter cell valve reflect not only the environmental conditions the parent cell was subjected to, but also the location of organelles within the parental cell (Shirokawa and Shimada 2016). As a result, diatoms exhibit phenotypic plasticity, and structural variation in shape and pattern of a cell are reflected during size diminution during the diatom life cycle. Size reduction may occur over many generations, and partial renewal inheritance occurs such that one valve is always a parental cell in contrast to the new cell generated (Shirokawa and Shimada 2016). Ensemble surface features could be measured for parent and daughter cells, and values for parent cells could be plotted and regressed on values for daughter cells to determine a regression coefficient. Potentially, hypotheses concerning cytoplasmic heritability (Shirokawa and Shimada 2016) could be devised and tested using ensemble surface measures.

Non-heritable factors could be studied using ensemble surface features. The combinations of such features that indicate buckling may be related to degree of phenotypic plasticity in diatom valve surfaces. Degree of buckling at the subcellular level may be influenced by environmental factors and indicative of degree of plasticity of a given diatom phenotype, and in turn, be indicative of the degree of phenotypic variation for a given diatom morphology.

### Phenotypic variation and ensemble surface features: epistasis and canalization

Diatom phenotype is affected by developmental changes. Such changes during development including valve formation may include epistasis and canalization (Waddington 1942). Phenotypic variation or the amount of variation in developmental factors is measurable as degree of canalization via the slope in a curved surface (Rice 1998). That is, minimum canalization (i.e., decanalization) is an increasing or maximum slope, while maximum canalization is a decreasing or minimum slope along a curve on a surface. Measurement of change phenotypically may be compared by the amount of developmental change as the phenotypic character becomes canalized (Rice 1998).

By contrast, epistasis represents distinct interactions among developmental states that are not additive. A phenotype gradient is the maximum slope of the curve on a surface and is comprised of first partial derivatives of a phenotype measure and are elements of the Jacobian. Eigenvectors of the Jacobian are phenotype gradients. An epistasis matrix has elements of second partial derivatives, and developmental epistasis is represented by the off-diagonal elements of the matrix, while the diagonal elements represent dominance and are eigenvalues (Rice 1998). This matrix is the Hessian, and the sum of the diagonal elements is the Laplacian. For additive effects given as zero valued off-diagonal elements and the same value for diagonal elements of the epistasis matrix, maximum slope is the same on a given surface and may be indicative of drift (Rice 1998). Ensemble surface measures of slopeness, peakedness and smoothness may be useful in constructing possible scenarios on the relation between morphogenesis, canalization and epistasis in diatom evolution.

## Conclusions

Quantifying diatom surface features from parametric 3D models enabled the analysis of multiple changes in morphology during a valve formation sequence. The analyses were centered on the combination of surface descriptors called ensemble surface features—slopeness, peakedness, smoothness, and flatness—as quantitative changes that could be compared among taxa. Three exemplars were used. *Actinoptychus senarius* was characterized by a combination of peakedness and smoothness, *Arachnoidiscus ehrenbergii* was characterized by a different combination of smoothness and peakedness, and *Cyclotella meneghiniana* was characterized by a varying combination of all the ensemble surface features measured.

For valve formation, the degree to which each taxon valve surface changed was measurable and elucidated for steps that were clustered and matched to Schmid and Volcani’s (1983) schema. Ensemble surface measures were used to determine that in stage one of valve formation, early change in the valve was horizontally flatter but sloped in terms of vertical changes. Midway through valve formation as stage two, horizontal and vertical surface changes increased in peakedness although smoothness of areas was also evident. Finally, toward completion of the valve at stage three, all ensemble surface measures contributed in varying degrees to the final surface “terrain” in terms of flat, smooth, sloped, and peaked features.

Matching ensemble surface features to definable stages in cytological processes, such as the position of organelles during mitosis or determining the degree to which ensemble surface features are a result of epistasis, is potentially in the offing and of interest in gaining an understanding of the role that cytology, inheritance and evolution have in diatom morphogenesis. Ensemble surface features have been demonstrated to be a quantifiable basis in morphological changes during diatom morphogenesis. Such features have potential application in studies of diatom valve formation with regard to phenotypic plasticity, epistasis and canalization.

